# VAP spatially stabilizes dendritic mitochondria to locally fuel synaptic plasticity

**DOI:** 10.1101/2023.01.16.524245

**Authors:** Ojasee Bapat, Tejas Purimetla, Sarah Kruessel, Christina Thum, Fiona Rupprecht, Monil Shah, Julian D. Langer, Vidhya Rangaraju

**Affiliations:** Max Planck Florida Institute for Neuroscience, Jupiter, FL 33458, USA; International Max Planck Research School for Synapses and Circuits, Jupiter, FL 33458, USA; Max Planck Institute for Brain Research, Frankfurt 60438, Germany; Max Planck Institute of Biophysics, Frankfurt 60438, Germany; Johns Hopkins University School of Medicine, Baltimore, MD 21205, USA; Thermo Fisher Diagnostics GmbH, Henningsdorf 16761, Germany

## Abstract

Synapses are pivotal sites of memory formation and undergo plasticity in response to external inputs. Consequently, synapses are hotspots of energy consumption and are susceptible to dysfunction when their energy supplies are perturbed. Mitochondria are stabilized near synapses via cytoskeletal tethering and serve as local energy supplies to fuel synaptic plasticity. However, the mechanisms that tether and stabilize neuronal mitochondria for long durations and determine the spatial dendritic segment supported during synaptic plasticity are unknown. We identified a list of novel mitochondrial-cytoskeletal interactors in neurons using APEX-based proximity labeling. We narrowed down the protein candidates that exclusively tether mitochondria to actin near postsynaptic spines using high-resolution Airyscan confocal imaging. We find that VAP, the vesicle-associated membrane protein-associated protein implicated in Amyotrophic Lateral Sclerosis and interacts with the endoplasmic reticulum, stabilizes mitochondria via actin near the spines. To test if the VAP-dependent stable mitochondrial compartments can locally support synaptic plasticity, we investigated individual spines stimulated by two-photon glutamate uncaging for spine plasticity induction and their adjacent spines. We find that, along with actin, VAP functions as a spatial stabilizer of mitochondrial compartments to sustain synaptic plasticity for up to ~60 min and as a spatial ruler that determines the ~30 μm length of the dendritic segment supporting synaptic plasticity.

## Introduction

Neurons have a complex morphology where most synapses are located far from the cell body. With increasing awareness of local processing at synapses (Holt et al., 2019; Nishiyama and Yasuda, 2015; Rangaraju et al., 2017), it is clear that synapses are hotspots of energy consumption. Therefore, a local energy source is necessary to cope with the fast and sustained energy demands of synapses (Jang et al., 2016; Li et al., 2004; Pathak et al., 2015; Rangaraju et al., 2014; Rangaraju et al., 2019a; Smith et al., 2016), as mere ATP diffusion from the cell body is insufficient (Hubley et al., 1996). Consistent with this notion, we recently showed that mitochondria form unique temporally (60 – 120 min) and spatially (30 μm) stable compartments by tethering to the cytoskeleton, actin and microtubule, in dendrites (Rangaraju et al., 2019a). The definition of a mitochondrial compartment, comprising single or multiple mitochondria, is where a sub-region of a compartment shows continuity with the rest of the compartment in fluorescence loss in photobleaching and photoactivation experiments ((Rangaraju et al., 2019a), see Methods). Notably, we found that perturbation of a dendritic mitochondrial compartment function abolishes protein synthesis-dependent plasticity in spines (Rangaraju et al., 2019a). Therefore, these stable mitochondrial compartments serve as local energy supplies to fuel synaptic plasticity in dendrites. What remains unknown is the mechanism that stabilizes mitochondria locally in dendrites and enables synaptic plasticity.

In contrast to dendritic mitochondria, most mitochondria are motile in axons within the long durations (minutes to hours) of plasticity formation and maintenance (Bradshaw et al., 2003; Kang and Schuman, 1996; Martin et al., 1997; Rangaraju et al., 2019a; Vickers et al., 2005). Interestingly, when the cytoskeleton (actin or microtubule) is depolymerized, stable dendritic mitochondria get destabilized, but in contrast, the motile axonal mitochondria get stabilized (Ligon and Steward, 2000; Rangaraju et al., 2019a). This observation suggests that dendritic mitochondria interact differently with the cytoskeleton than axonal mitochondria. This mechanism might be facilitated by exclusive protein(s) tethering mitochondria to the cytoskeleton in dendrites that are either absent in axons or not associated with axonal mitochondria.

Many proteins and their mechanisms involved in docking axonal mitochondria have been well studied near axonal terminals and during axon branching (notably, most of these studies have focused on mitochondrial-microtubule interactions (Courchet et al., 2013; Kang et al., 2008; MacAskill et al., 2009; Wang and Schwarz, 2009) and a few on mitochondrial-actin interactions (Basu et al., 2021; Spillane et al., 2013)). However, the mechanisms required for stabilizing dendritic mitochondria near postsynaptic spines, particularly to support synaptic plasticity, remain largely unexplored. Given that dendritic and axonal compartments are biochemically, structurally, and functionally distinct, and the mitochondria in these two compartments also behave very differently (Lewis et al., 2018; Popov et al., 2005; Rangaraju et al., 2019a; Rangaraju et al., 2019b; Shepherd and Harris, 1998), it is important to characterize the specialized mitochondrial mechanisms in dendrites and how they enable synaptic plasticity.

Here, we set out to find the protein tether required for mitochondrial-cytoskeletal interaction and if it determines the spatial size of the dendritic segment supported during synaptic plasticity. For instance, the length of the mitochondrial compartment (30 μm) might determine the spatial spread and availability of ATP within a dendritic segment. Such a spatial organization might be necessary for clustered synaptic plasticity, where a cluster of neighboring spines within a certain distance from a plasticity-induced spine exhibit plasticity (Govindarajan et al., 2011; Govindarajan et al., 2006). As clustered plasticity is important during learning, experience, and development (Fu et al., 2012; Kleindienst et al., 2011; Makino and Malinow, 2011; McBride et al., 2008; Takahashi et al., 2012; Wilson et al., 2016), it is essential to understand the underlying molecular mechanisms that support them.

Using advances in proximity-labeled mitochondrial proteome labeling and imaging mitochondrial-actin tethering and stabilization (Hung et al., 2017; Lam et al., 2015; Schiavon et al., 2020), we have identified novel proteins tethering mitochondria to actin exclusively in dendrites. In addition, we find that the vesicle-associated membrane protein-associated protein (VAP) (Gomez-Suaga et al., 2019; Guillen-Samander and De Camilli, 2022; Teuling et al., 2007) plays a central role in stabilizing dendritic mitochondria for long durations of synaptic plasticity formation and maintenance, and determining the spatial segment supported during synaptic plasticity.

## Results

### Proximity labeling strategy to label proteins interacting with mitochondria

To identify the proteins spatially stabilizing dendritic mitochondria, we first screened for proteins required for mitochondrial-cytoskeletal interaction in dendrites. We employed a proximity labeling strategy based on the enzyme APEX2, an engineered form of a soybean ascorbate peroxidase (Rhee et al., 2013). APEX2 can catalyze the promiscuous biotinylation of proteins within a few nanometers of its active site (Rhee et al., 2013). Most studies with APEX2 have been done in either non-neuronal cells or non-hippocampal neurons. As we are interested in studying synaptic plasticity in hippocampal dendrites, we first validated and characterized APEX2 biotinylation of proteins in dendritic compartments of primary hippocampal neuronal cultures. We expressed APEX2 either on the outer mitochondrial membrane (APEX-OMM, APEX2 is simplified as APEX in the rest of the text) or in the mitochondrial matrix (APEX-matrix) and labeled the neurons with biotin phenol and hydrogen peroxide (Fig. 1A, See Methods) (Hung et al., 2017; Rhee et al., 2013). APEX-OMM expression colocalized with the spatially confined mitochondrial signal (visualized by EGFP targeted to mitochondria) in neurons (Fig. S1A and B). However, compared to the spatially confined expression of APEX-OMM, the biotinylation signal of APEX-OMM was cytosolically distributed, indicating the diffusion of the biotinylated proteins from the mitochondrial membrane surface to the neighboring neuronal regions (Fig. 1B, D). In contrast, APEX-Matrix expressing neurons exhibited a spatially-confined biotin signal overlapping with the spatially confined expression of APEX-Matrix, indicating the compartmentalization of biotinylated proteins within the mitochondrial matrix (Fig. 1C, E). Furthermore, biotin labeling was not observed in the absence of biotin phenol or the presence of an alternative substrate, biotin (Fig. S1C and D), suggesting that biotinylation of proteins is specific to the presence of biotin phenol as the substrate. These experiments confirmed that APEX-OMM-based proximity labeling could be used to biotinylate proteins present on and interacting with the mitochondrial outer membrane in hippocampal neurons.

**Figure 1.**
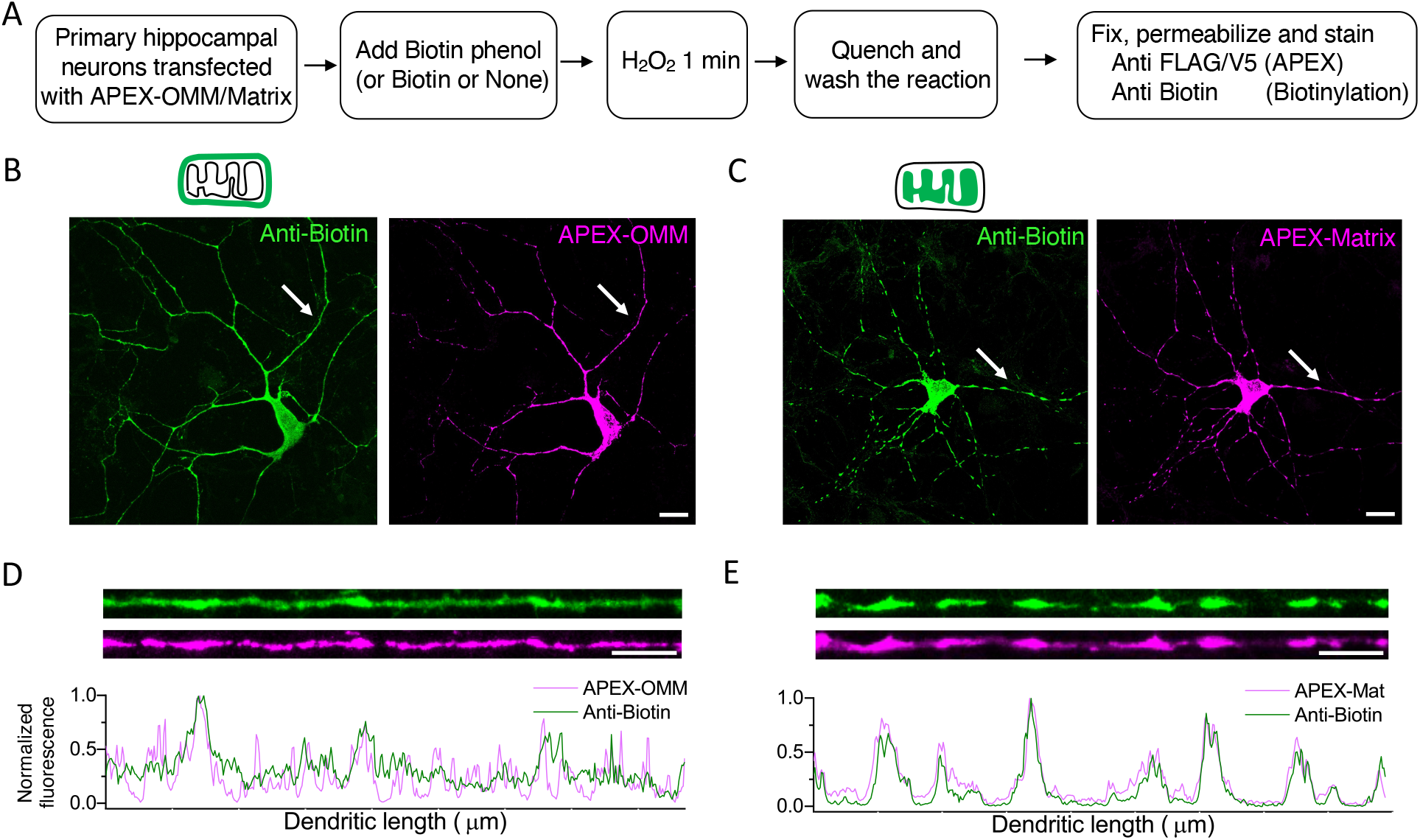
APEX-strategy to label proteins interacting with neuronal mitochondria. (A) Experimental workflow for APEX-based proteome labeling of the mitochondrial outer membrane (OMM) and mitochondrial matrix. Representative images showing the biotinylated proteome (left, green) and APEX-expression (right, magenta) on the OMM (B) and within the matrix (C). Scale bar, 20 μm. Line profiles of the respective dendrites pointed in B and C (white arrows) show diffused biotin labeling with APEX-OMM (D) but spatially confined biotin labeling with APEX-Matrix (E) in comparison to APEX-OMM and APEX-Matrix expression, respectively. Scale bar 10 μm. See also Figure S1.

### Identifying proteins interacting with the outer mitochondrial membrane and actin

To isolate proteins interacting with the mitochondrial outer membrane, we expressed primary hippocampal neuronal cultures with APEX-OMM, labeled the neurons with biotin phenol and hydrogen peroxide, enriched the biotinylated proteins using streptavidin, digested and analyzed them using liquid chromatography-coupled tandem mass spectrometry (LC-MS) (Fig. 2A). Neurons labeled either in the absence of APEX-OMM or hydrogen peroxide were used as negative controls (Fig. 2B). Western blot analysis of the streptavidin-enriched proteins showed biotinylated proteins in APEX-OMM samples compared to the negative control without APEX-OMM (Fig. S2A); with an equal amount of total protein loaded in both conditions confirmed by Coomassie stain (Fig. S2B).

**Figure 2.**
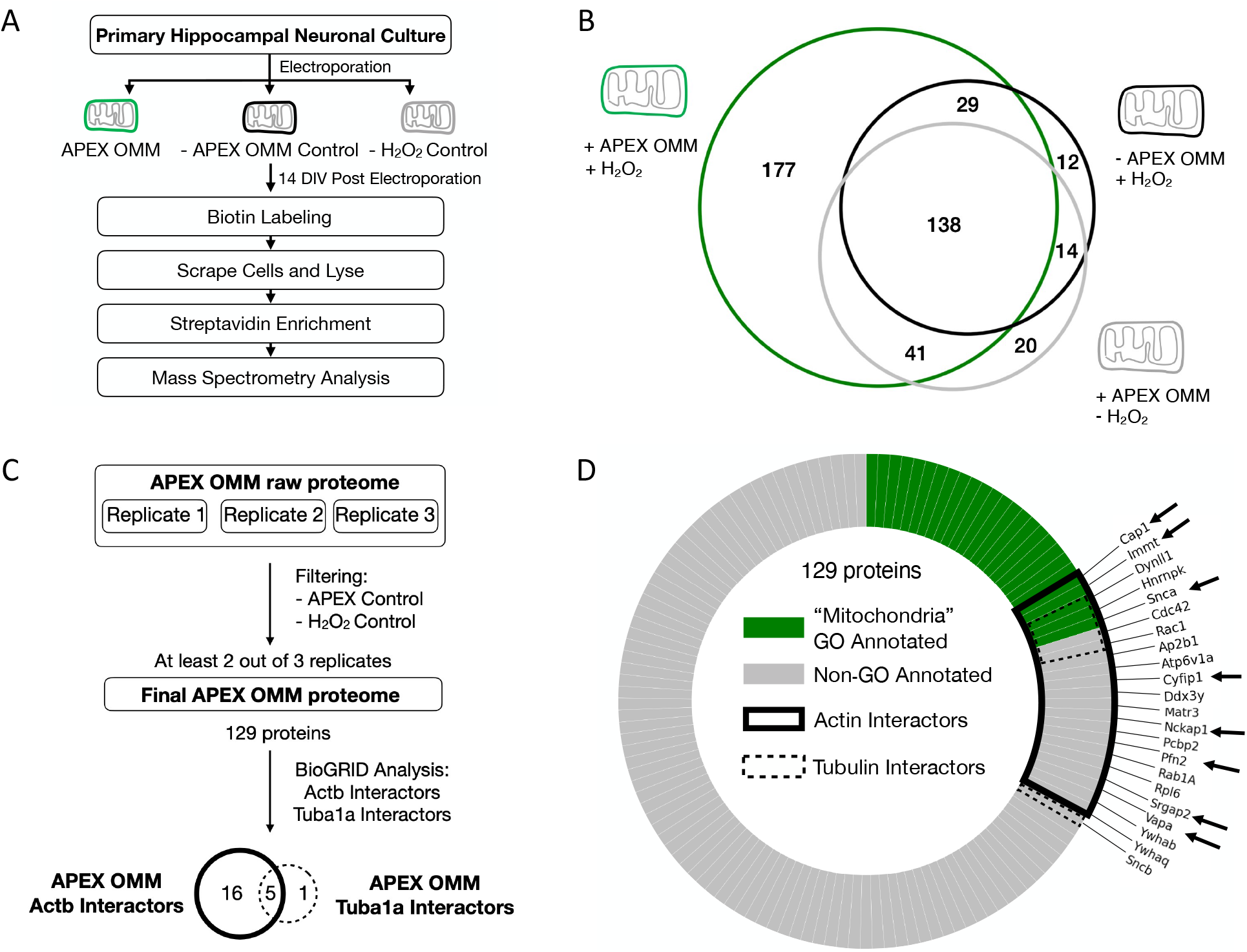
Identifying novel actin interacting proteins on the outer mitochondrial membrane. (A) Experimental workflow for APEX-OMM proteome labeling in primary hippocampal neuronal cultures for mass spectrometry analysis. (B) Representative replicate showing the OMM proteome yield (green) after subtracting the proteins measured in controls –in the absence of APEX-OMM (black) and the absence of hydrogen peroxide (gray). The proteins detected in controls are endogenous biotinylated proteins (pyruvate carboxylase, 3-methylcrotonyl CoA carboxylase, propionyl CoA carboxylase, and acetyl CoA carboxylase) and non-specific proteins binding to streptavidin beads during enrichment. As we followed a stringent criterion of excluding the proteins found in controls from experimental samples, irrespective of their intensities (see Methods), there might be an overestimation of proteins in the two control samples. (C) Flowchart of the mass spectrometry proteome analysis of three replicates, following filtering for controls as in B, yielding 129 proteins from two out of three replicates and identifying actin (Actb) and tubulin (Tuba1a) interactors on the OMM using BioGRID. (D) 129 proteins of the OMM proteome analyzed for gene ontology (GO) annotation for the term “Mitochondria” (“Mitochondria GO annotated, green), and proteins not GO annotated as “Mitochondria” (Non-GO annotated, gray). 21 proteins identified in BioGRID as actin interactors (Actin interactors, black line), 6 proteins identified in BioGRID as tubulin interactors (Tubulin interactors, dotted black line), and the 8 actin-interacting OMM proteins selected for the next round of screening (black arrows). See also Figure S2, Table S1, and S2.

An exclusive APEX-OMM protein list was obtained from three replicates after subtracting proteins found in negative controls without APEX-OMM and hydrogen peroxide (Fig. 2B, representative replicate, Table S1). Next, we consolidated a list of exclusive APEX-OMM proteins found in at least two of the three replicates resulting in a total of 129 proteins (Fig. 2C, Table S1). These 129 proteins comprise proteins on the mitochondrial membrane (26 proteins with ‘Mitochondria’ GO annotation, Fig. 2D, green) and proteins interacting with the mitochondrial membrane (123 proteins without ‘Mitochondria’ GO annotation, Fig. 2D, gray). We further analyzed the 129 proteins using a curated repository for protein interactions, BioGRID (Oughtred et al., 2021), to identify known protein interactors of actin and tubulin (see Methods). This analysis resulted in 16 exclusive actin-interacting proteins, 1 exclusive tubulin-interacting protein, and 5 proteins that were actin and tubulin interactors (Fig. 2C, D, Table S2). From these total 21 proteins, we focused on the following 8 proteins – CAP1, IMMT, SNCA, CYFIP1, NCKAP1, PFN2, SRGAP2, and VAPA, for further analysis (Fig. 2D, Table S2, green). Most proteins with multiple roles in basic cellular function and signaling, such as RAB1A, RAC1, RPL6, YWHAB, YWHAQ, etc., were not followed up for this study (Fig. 2D).

### VAP, including other proteins, is required for mitochondria-actin tethering in dendrites

We then determined the requirement of the 8 protein candidates in tethering mitochondria to actin in dendrites. We genetically knocked down each candidate protein and measured the mitochondrial-actin interaction efficiency in dendrites and axons. For this purpose, we established a method based on Lifeact-GFP (Riedl et al., 2008), an actin-visualization marker. Lifeact-GFP has an actin-interacting domain and can be targeted to the OMM (Fis1-Lifeact-GFP)(Schiavon et al., 2020). At reduced expression levels of Lifeact-GFP in the OMM, only the Lifeact-GFP that binds and accumulates near actin, and is therefore immobile, is detectable by fluorescence (Fig. 3A). In contrast, the low fraction Lifeact-GFP that freely diffuses on the OMM, and is mobile, is undetectable. To image Fis1-Lifeact-GFP, we employed high-resolution Airyscan confocal that revealed the fraction of the OMM area colocalizing with actin, which we measure as a proxy for mitochondria-actin interaction (Fig. 3, see Methods). The average mitochondria-actin interaction was 55 ± 2 % in control dendrites (Fig. 3C, see Methods). Knocking down the 8 protein candidates, CAP1, IMMT, SNCA, CYFIP1, NCKAP1, PFN2, SRGAP2, and VAPA reduced the mitochondria-actin interaction in dendrites (Fig. 3B, C), suggesting their significance in tethering mitochondria to actin.

**Figure 3.**
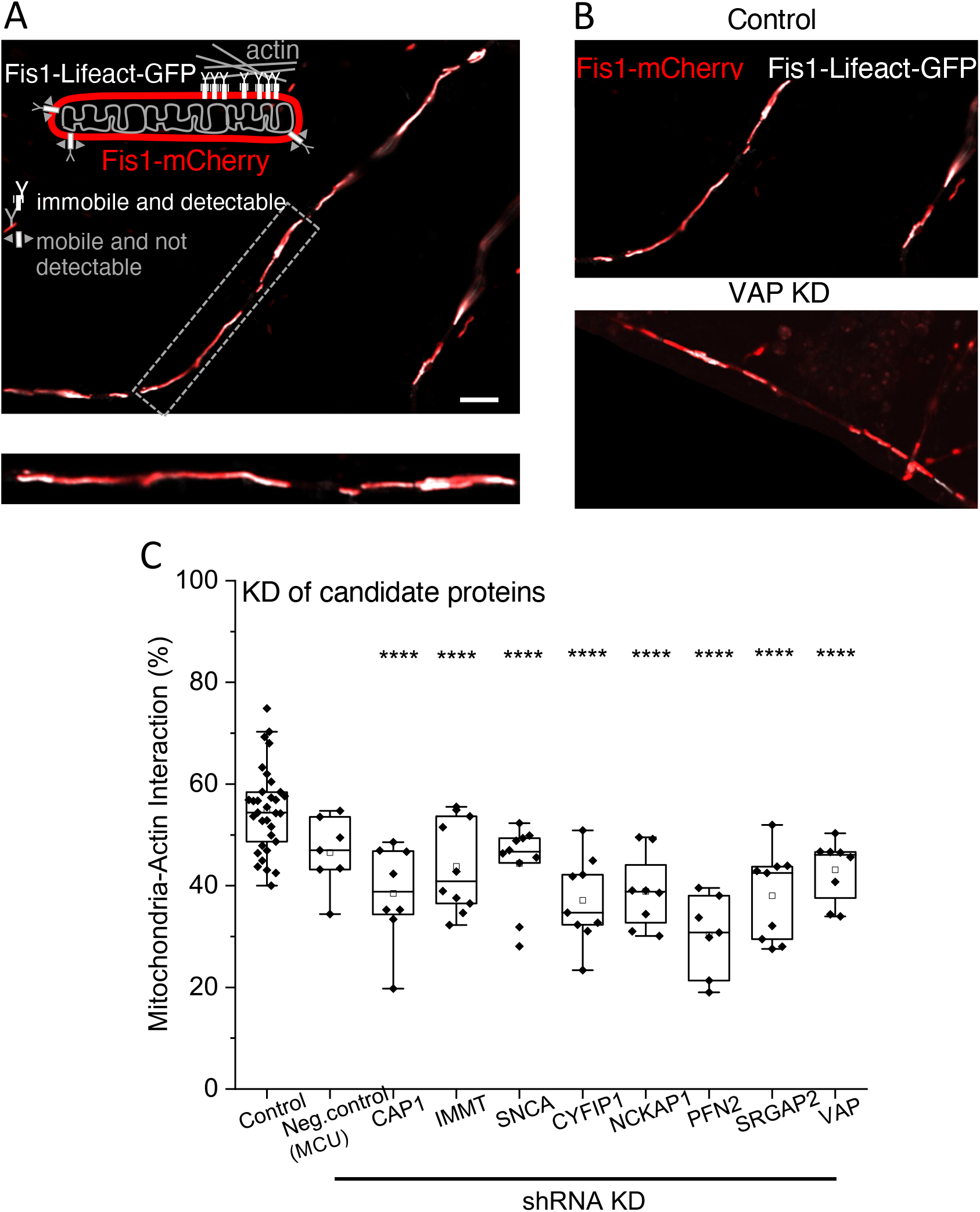
Identifying novel proteins required for mitochondria-actin tethering in dendrites. (A) Representative image of a neuronal dendrite transfected with Fis1-mCherry (red) and Fis1-Lifeact-GFP (white) showing mitochondrial regions interacting with actin (white). Top left inset: cartoon illustrating the labeling strategy used for visualizing mitochondria and mitochondria-actin interaction regions. The Gray dashed box depicts the dendritic segment magnified for better visualization (bottom inset). Scale bar, 5 μm. (B) Representative images showing fewer mitochondrial (red) regions interacting with actin (white) in VAP KD dendrites (bottom) compared to control dendrites expressing control shRNA (top). (C) Significant reduction in the average mitochondria-actin interaction percentage in all 8 protein knockdown conditions compared to control shRNA-expressing dendrites. The mitochondrial uniporter (MCU) was used as a negative control, given its localization in the mitochondrial inner membrane and the absence of any known actin association. Control (control shRNA), Neg. control (MCU shRNA as negative control), CAP1, IMMT, SNCA, CYFIP1, NCKAP1, PFN2, SRGAP2, VAP. n ≥ 7 neurons from ≥ 2 animals. Oneway ANOVA, Tukey test, ****p ≤ 0.000001. See also Figure S3.

As a positive control, neurons were treated with actin depolymerizing agents, Cytochalasin (Cyto-D) and Latrunculin-B (Lat-B), to confirm the ability of Fis1-Lifeact-GFP to detect a reduction in mitochondria-actin interaction (Fig. S3A). As a negative control for the Fis1-Lifeact-GFP measurements, neurons were treated with a microtubule depolymerizing agent that does not affect actin, Nocodazole (Noco). Noco treatment did not affect mitochondria-actin interaction (Fig. S3A). In addition, the expression of Fis1-Lifeact-GFP did not cause any abnormal aggregation of actin in dendrites (Fig. S3B).

Furthermore, axonal mitochondrial interaction with actin was unaffected when the 8 protein candidates were knocked down (Fig. S3 C, E). This data suggests that the identified 8 protein candidates are exclusive to mitochondrial-actin tethering in dendrites and not axons. Furthermore, when treated with actin depolymerizing agents, mitochondrial-actin interaction in axons was unaffected (Fig. S3D). These data are consistent with our previous observations that axonal mitochondria stabilize on cytoskeletal depolymerization while dendritic mitochondria destabilize (Ligon and Steward, 2000; Rangaraju et al., 2019a), suggesting that different mechanisms tether mitochondria in axons compared to dendrites (see Discussion).

### VAP stabilizes mitochondrial compartments in dendrites

The 8 proteins tethering mitochondria to actin might not necessarily stabilize mitochondrial compartments in dendrites. Therefore, we further refined the 8 protein-candidate list by identifying the protein(s) required for stabilizing mitochondria. Using our previously established photoactivation method to quantify mitochondrial stability (Rangaraju et al., 2019a) (Fig. 4A), we knocked down 3 of the 8 protein candidates (VAP, SNCA, SRGAP2) for this study and measured mitochondrial compartment size and stability in dendrites. Knocking down VAP, SNCA, and SRGAP2 shortened the mitochondrial compartment length (Fig. S4 A-F, see Methods). We then measured mitochondrial stability in VAP, SNCA, and SRGAP2 KD dendrites by monitoring the photoactivated compartment fluorescence over time and measuring the compartment stability index (Fig. 4B, C, see Methods). Mitochondria in control dendrites showed a modest decrease in photoactivated compartment fluorescence 60 minutes post-photoactivation, perhaps corresponding to fluorescent protein leak from mitochondria during basal-level mitochondrial dynamics such as fission and fusion. Meanwhile, dendrites lacking VAP showed a dramatic decrease in photoactivated compartment fluorescence 60 minutes post-photoactivation, corresponding to destabilized mitochondrial compartments (Fig. 4A-C). This result is similar to the mitochondrial compartment length shortening and destabilization observed on actin perturbation (Rangaraju et al., 2019a), suggesting VAP’s role in actin-tethering and mitochondrial stabilization in dendrites. While knocking down SNCA and SRGAP2 showed a moderate destabilization of mitochondrial compartments, it was not statistically significant (Fig. 4-C).

**Figure 4.**
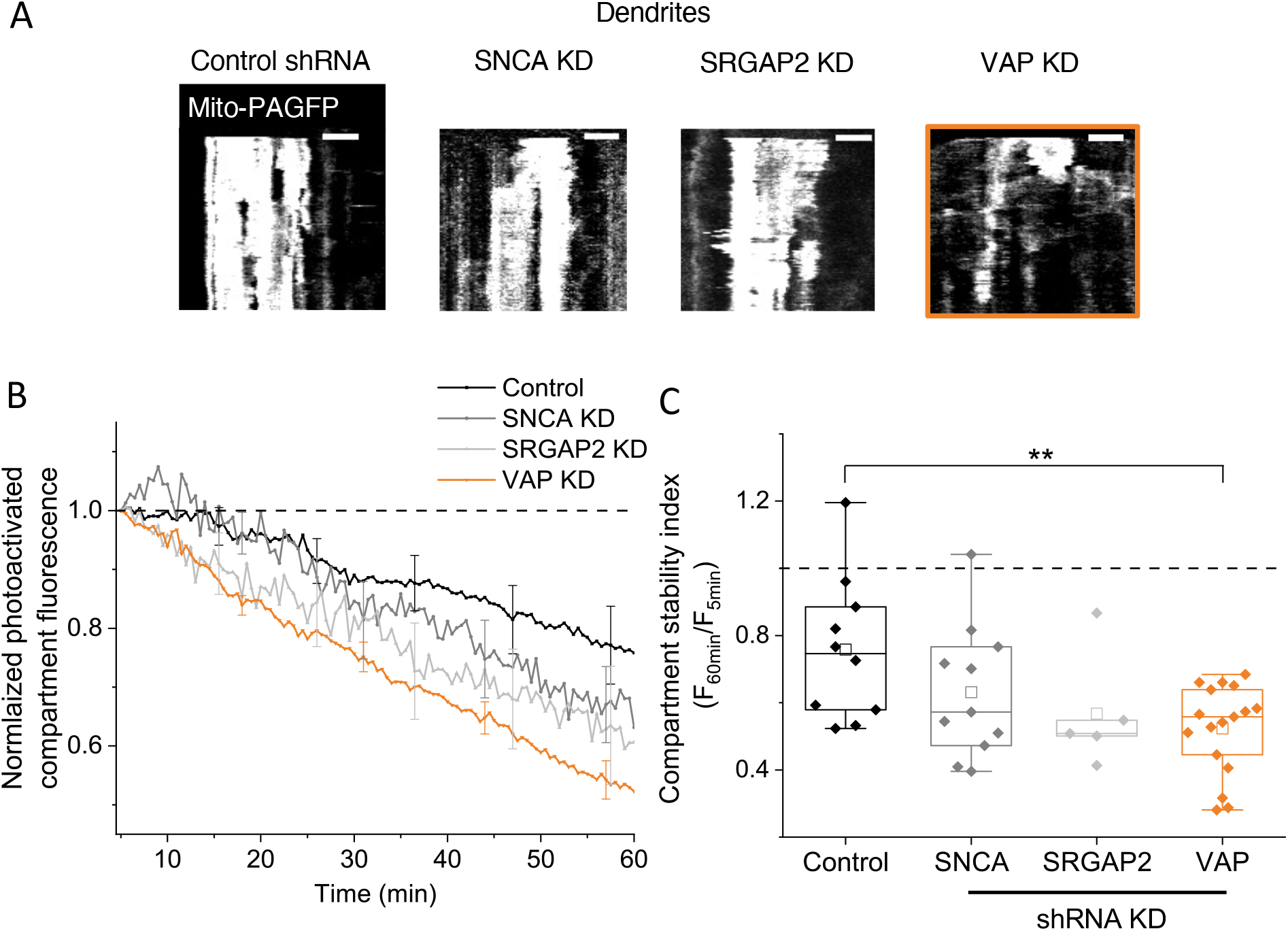
VAP spatially stabilizes mitochondria in dendrites. (A) Representative kymographs of photoactivated mitochondrial compartments show stable mitochondrial compartments in control shRNA-expressing dendrite (left); shortened and modest destabilization of mitochondrial compartments in SNCA KD and SRGAP2 KD dendrites (middle); and shortened and destabilized mitochondrial compartments in VAP KD dendrites (right, orange box). Scale bar, 5 μm. (B) The average time course of photoactivated mitochondrial compartment fluorescence shows a modest decrease in control shRNA expressing dendrites (control), a moderate decrease that was not statistically significant in SNCA KD (gray) and SRGAP2 KD (light gray) dendrites, and a dramatic and statistically significant decrease in VAP KD dendrites (orange). (C) The mitochondrial compartment stability index showed a statistically significant decrease in VAP KD dendrites compared to control, SNCA KD, and SRGAP2 KD dendrites. n ≥ 4 neurons from ≥ 2 animals. One-way ANOVA, Tukey test, **p ≤ 0.01. See also Figure S4.

### VAPB is enriched near mitochondria in dendrites but not in axons

VAP is an endoplasmic reticulum (ER)-associated protein that tethers ER with other organelles such as endosomes and mitochondria, and its dysfunction is implicated in Parkinson’s Disease (PD), Amyotrophic Lateral Sclerosis (ALS), and frontotemporal dementia (Dong et al., 2016; Gomez-Suaga et al., 2019; Guillen-Samander and De Camilli, 2022; Lindhout et al., 2019; Teuling et al., 2007). VAP is present in two isoforms, VAPA and VAPB (Guillen-Samander and De Camilli, 2022). As one isoform can compensate for the absence of the other, we deleted both isoforms for our gene deletion experiments. However, the local association of each of these isoforms, VAPA and VAPB, with mitochondria is unclear in neuronal compartments.

We, therefore, visualized endogenous VAPA and VAPB by immunostaining and measured its association (Pearson correlation coefficient, see Methods) with mitochondria in both dendrites and axons in fixed neurons. VAPA (A, immunodetection) is enriched near dendritic and axonal mitochondria (Fig. 5A, D). However, VAPB (B, immunodetection) is only enriched near dendritic mitochondria and not axonal mitochondria (Fig. 5B-D). Visualization of VAPB in live neurons (B_live_, VAPB-emerald) also confirmed its enrichment near dendritic mitochondria and not axonal mitochondria (Fig. 5D, S5). The enrichment of VAPB only near dendritic mitochondria, and its low expression compared to VAPA, could be why we did not detect it in our APEX-OMM proteome (see Discussion). Our results suggest that the VAPB isoform, exclusively enriched near dendritic mitochondria, might determine the dendrite-specific role of VAP in stabilizing mitochondria.

**Figure 5.**
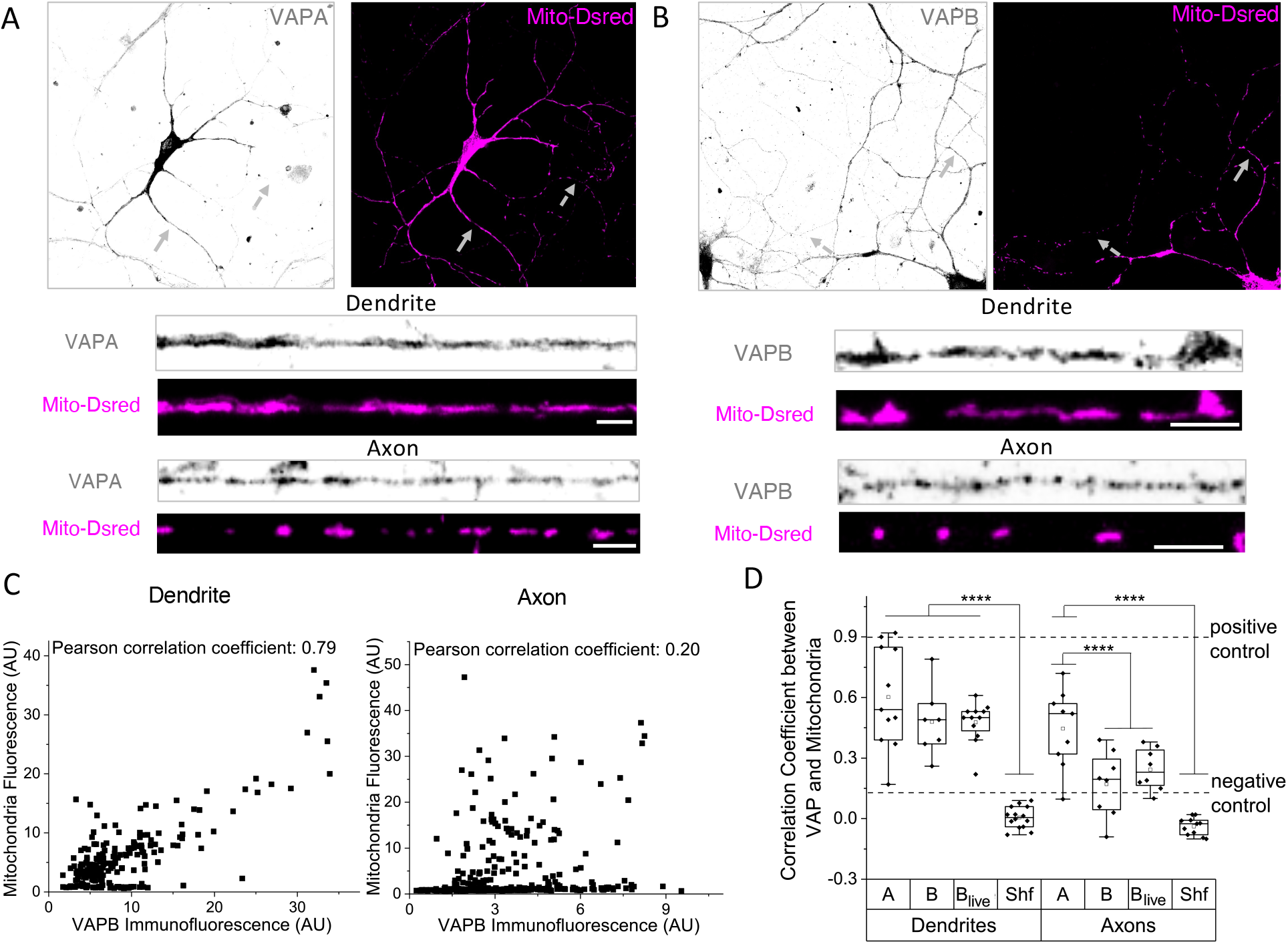
VAPB is enriched near mitochondria in dendrites but not in axons. (A) Representative images of neurons immunostained for VAPA (left, gray) and mitochondria (right, Mito-DsRed, magenta). Arrows depicting dendrite (gray) and axon (dotted gray) magnified for better visualization (bottom insets), showing local enrichment of VAPA near dendritic and axonal mitochondria. VAPA antibody was validated in VAPA knocked down neurons. Scale bar, 5 μm. (B) Representative images of neurons immunostained for VAPB (left, gray) and mitochondria (right, Mito-DsRed, magenta). Arrows depicting dendrite (gray) and axon (dotted gray) magnified for better visualization (bottom insets), showing local enrichment of VAPB near dendritic mitochondria but not near axonal mitochondria. VAPB antibody was validated in VAPB knocked down neurons. Scale bar, 5 μm. (C) Representative correlation between individual fluorescent pixel intensities of mitochondria (Mito-DsRed) and VAPB show a strong correlation in the dendrite (left) but not in the axon (right). (D) The average correlation coefficient measured shows a strong correlation between mitochondria and VAPA (A), VAPB (B) in fixed dendrites, and VAPB in live dendrites (B_live_, VAPB-emerald), compared to shuffled control (Shf, see Methods). In axons, a strong correlation between mitochondria and A was observed, but not between mitochondria and B and B_live_. n ≥ 8 neurons from ≥ 2 animals. One-way ANOVA, Tukey test, ****p ≤ 0.000001. As a positive control, the correlation between two different fluorescent tags (Mito-DsRed and Mito-PAGFP) targeted to the mitochondrial matrix was used, which provided the upper limit of the correlation coefficient. As a negative control, the correlation between a mitochondrial (Mito-DsRed) and a non-mitochondrial tag was used, which provided the lower limit of the correlation coefficient. See also Figure S5.

### VAP serves as a spatial ruler determining the dendritic segment fueled by mitochondria during synaptic plasticity

Next, we determined if VAP-dependent mitochondrial stabilization can locally support synaptic plasticity. Mitochondrial compartments are stable for ~60 min and are essential to fuel protein synthesis-dependent synaptic plasticity within 30 μm dendritic segments (Rangaraju et al., 2019a). So, we hypothesized that if VAP is required for the temporal and spatial stabilization of dendritic mitochondria, the absence of VAP will affect both the temporal sustenance of synaptic plasticity formation and the spatial dendritic segment exhibiting synaptic plasticity (Fig. 6F). We induced single-spine plasticity using two-photon glutamate uncaging (0.5 Hz,120 pulses) in PSD95- or Homer2-positive spines in the presence of forskolin to activate the PKA pathway to induce long-term structural plasticity (Govindarajan et al., 2011) (Fig. 6A). We then measured spine size increase in the stimulated spine and adjacent spines in the same dendritic segment (Fig. 6). We also measured spine size from unstimulated spines in sister dendrites as control (Fig. 6C).

**Figure 6.**
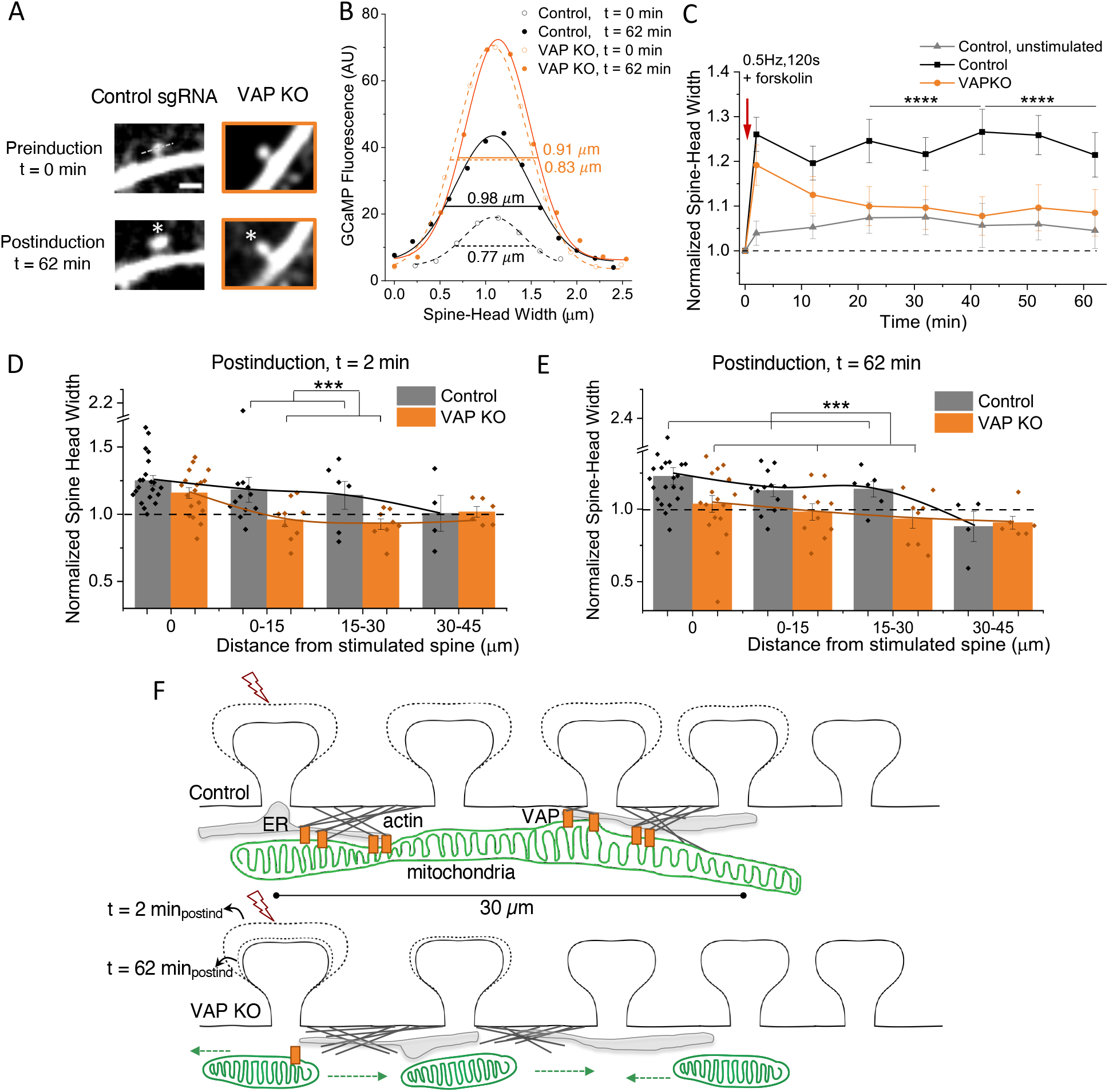
VAP spatially stabilizes mitochondria to locally fuel synaptic plasticity. (A) Representative plasticity-induced spine (white asterisk) measured at 0 min (preinduction) and 62 min (postinduction) showed an increase in spine-head width in control neurons expressing control sgRNA (Left, Control), but not in VAP KO neurons (Right, VAP KO, orange box). Scale bar, 2 μm. (B) GCaMP fluorescence intensity along the line crossing the center of the spine-head (gray dashed line in A), fit to a Gaussian to measure the full width at the half maxima, the spinehead width (see Methods), in control spines expressing control sgRNA (black) and VAP KO spines (orange), before (empty circles, t=0 min) and after plasticity induction (full circles, t=62 min). (C) The average time course showed a concomitant increase in spine-head width in control spines expressing control sgRNA (black) but not in VAP KO spines (orange) at 62 min post-spine plasticity induction. Control, unstimulated spines from sister dendrites, also exposed to forskolin, did not exhibit structural plasticity (gray). n ≥ 19 spines from ≥ 5 animals. One-way ANOVA, Tukey test, ****p ≤ 0.000001. (D, E) Histogram showing concomitant increase in spine-head width 2 min and 62 min post-plasticity induction in control spines expressing control sgRNA within 0 – 30 μm from plasticity-induced spines (gray), but not in spines within 30-45 μm distance. In VAP KO neurons (orange), 2 min post-plasticity induction (D), an increase in spine-head width was observed in plasticity-induced spines (orange) but not in adjacent spines within 0-45 μm, whereas spine plasticity was not observed in both plasticity-induced and adjacent spines 62 min post-plasticity induction (E). n ≥ 4 neurons with up to 3 spines each from ≥ 2 animals. One-way ANOVA, Tukey test, ***p ≤ 0.001. (F) Illustration showing the significance of VAP as a spatial organizer in stabilizing mitochondrial compartments to fuel plasticity-induced spines (lightning bolt) and adjacent spines. Top: control dendritic segment with normal VAP expression and mitochondrial stabilization exhibits spine structural plasticity in stimulated spines and adjacent spines within 30 μm. Bottom: VAP deletion shortens and destabilizes mitochondria (short green mitochondria that are mobile with green dashed arrows) in the dendrites and abolishes the sustenance of spine structural plasticity in both stimulated and adjacent spines within 30 μm. VAP (orange full rectangles) serves as a spatial organizer in stabilizing mitochondrial compartments (green mitochondria) via ER-actin-tethering (gray ER and gray actin filaments) for sustained synaptic plasticity formation. VAP also functions as the spatial ruler determining the 30 μm dendritic segment fueled by a mitochondrial compartment. These data emphasize the importance of VAP as a molecular tether in locally stabilizing a mitochondrial compartment to fuel the synaptic and clustered synaptic plasticity required for learning and development in a spatially confined manner in dendrites.

VAP depletion showed a modest and negligible increase in spine size at baseline (Fig. S6A) and did not affect spine density (Fig. S6B) compared to control neurons. VAP depletion also did not affect spine calcium influx in plasticity-induced spines compared to control neurons (Fig. S6C and D).

The absence of VAP reduced spine morphological plasticity in the stimulated spine, indicating the importance of stable mitochondrial compartments at the base of the plasticity-induced spine (Fig. 6A-C). Interestingly, VAP deletion did not affect the early stages of synaptic plasticity induction as the measured spine-head width increase was similar to that of control spines at t=0 and 12 min (Fig. 6C). However, post-plasticity induction over longer durations of t=22 to 62 min, spines lacking VAP were unable to sustain the structural plasticity. This data suggests the importance of stabilized mitochondrial compartments in supporting the long durations of synaptic plasticity formation and maintenance.

Furthermore, we investigated the ability of the spines adjacent to the plasticity-induced spine to exhibit structural plasticity, a phenomenon otherwise known as clustered synaptic plasticity. In control neurons, spines adjacent to the plasticity-induced spine within 0-15 μm showed structural plasticity consistent with earlier observations (Fu et al., 2012; Govindarajan et al., 2011; Kleindienst et al., 2011; Makino and Malinow, 2011; McBride et al., 2008; Takahashi et al., 2012)), with a similar trend in spines within 15-30 μm distance, and no apparent structural plasticity in spines within 30-45 μm distance, at t=2 and 62 min post-plasticity induction (Fig. 6D and E, gray bars, S6E). However, on VAP deletion, the ability of the adjacent spines to exhibit structural plasticity was affected as early as t=2 minutes post-plasticity induction (Fig. 6D and E, orange bars, S6E). These data suggest that the mitochondrial compartment length shortening and destabilization in the absence of VAP affect the long-term plasticity formation in stimulated and adjacent spines within a 30 μm distance. In summary, these results indicate that VAP functions as a spatial stabilizer that temporally sustains synaptic plasticity for up to ~60 min and as a spatial ruler that determines the ~30 μm spatial dendritic segment fueled by mitochondria during synaptic plasticity (Fig. 6F).

## Discussion

Mitochondrial dysfunction is associated with various disease states. As distal synapses rely on local energy supplies such as mitochondria, even brief interruptions in their local availability can result in long-lasting changes in synaptic and network properties, as observed in neurodegenerative diseases including Alzheimer’s (AD), PD, and ALS (Lee et al., 2018; Rangaraju et al., 2019b; Wang et al., 2019). Using recent advances in subcellular proteomic labeling, high-resolution imaging techniques to quantify mitochondrial-actin interactions, and stimulation of individual spines using two-photon glutamate uncaging for measurements of synaptic plasticity, we probe various mechanisms determining the spatial organization of dendritic mitochondria and its role in powering synaptic plasticity. We demonstrate the following: (1) identification of novel mitochondrial-actin protein interactors in neurons; (2) identification of novel mitochondrial-actin tethering proteins that are exclusive to neuronal dendritic compartments; (3) novel role of VAP in spatially stabilizing mitochondria in dendrites; (4) the VAPB isoform of VAP is exclusively enriched near dendritic mitochondria, and not near axonal mitochondria; (5) gene deletion of VAP disrupts sustained spine morphological plasticity in stimulated spines and adjacent spines within 30 μm; (6) VAP serves as the spatial stabilizer that determines the temporal sustenance of synaptic plasticity for 60 min and a spatial ruler that determines the 30 μm dendritic segment fueled by mitochondria during synaptic plasticity.

Using APEX-based proximity labeling, we identified 129 proteins in the neuronal OMM proteome that includes proteins present on the OMM and interacting with the OMM. Of these 129 proteins, 69 proteins (53%) overlap with the OMM proteome of HEK cells (Hung et al., 2017), which includes the 7 of the 8 mitochondrial-actin interacting proteins we investigated in our experiments (CAP1, SNCA, CYFIP1, NCKAP1, PFN2, SRGAP2, VAP, except for IMMT) (Table S1). The rest of the proteins we identified in the OMM proteome might be proteins specific to neurons. The proteins that were found in the HEK cell OMM proteome but not in our proteome might either be HEK cell-specific proteins, proteins undetected due to different mass spectrometry analysis strategies, or low expressing proteins that could not be detected in neurons (as neurons being non-dividing cells generally provide lower proteome yield compared to HEK cells). Furthermore, the BioGRID database we used to filter for actin-interacting proteins is constantly updated; therefore, not all actin interactors are fully annotated or identified. Our mitochondrial-actin interactor list will accordingly be updated with the growing literature in the future.

Dendritic mitochondria differ from axonal mitochondria in size, shape, and motility. Dendritic mitochondria can be as long as 30 μm in length, where multiple mitochondrial filaments stack against each other to make one physical compartment (Lewis et al., 2018; Popov et al., 2005; Rangaraju et al., 2019a). Dendritic mitochondria are also spatially stable for 60 to 120 min and during development (Faits et al., 2016; Rangaraju et al., 2019a). On the other hand, axonal mitochondria are short, between 0.5 to 2 μm in length, they are not stacked, and are motile within the same duration of 60 to 120 min (Lewis et al., 2018; Rangaraju et al., 2019a; Shepherd and Harris, 1998; Wang and Schwarz, 2009). However, it has been shown that mitochondria reach stable positions during axonal maturation and axonal branching (Courchet et al., 2013; Lewis et al., 2016; Spillane et al., 2013). In addition, the mechanisms by which dendritic and axonal mitochondria interact with the cytoskeleton are also strikingly different. When the cytoskeleton (actin or microtubule) is perturbed, stable dendritic mitochondria get destabilized, but in contrast, the motile axonal mitochondria get stabilized (Ligon and Steward, 2000; Rangaraju et al., 2019a). This observation suggests that dendritic mitochondria interact differently with the cytoskeleton than axonal mitochondria.

Interestingly, the 8 proteins we characterized as essential for mitochondrial-actin tethering in dendrites did not affect mitochondrial-actin tethering in axons when knocked down. This result suggests that mitochondria have different tethering and stabilization rules in axons compared to dendrites. For example, syntaphilin, which facilitates mitochondrial docking via the microtubule, is an axon-specific protein and is not present in dendrites (Kang et al., 2008). Similarly, FHL2 was recently shown to anchor mitochondria to actin at high glucose concentrations in axonal compartments (Basu et al., 2021). While this study did not investigate FHL2 in dendrites, knocking down FHL2 in neurons also affects mitochondrial-actin interaction in dendrites (our unpublished data), emphasizing that it may play a dual role in both neuronal compartments. Future studies should therefore identify more proteins that play a specific or unifying role in mitochondrial cytoskeletal tethering and stabilization in dendritic and axonal compartments. Furthermore, the current study only focused on mitochondrial-actin tethering and stabilization proteins in dendrites. Therefore, the mitochondria-tubulin interacting proteins and their role in dendritic mitochondrial tethering and stabilization should be investigated in future studies.

VAP is an ER resident protein found in membrane contact sites with various organelles such as Golgi, mitochondria, peroxisomes, endolysosomes, and plasma membrane. It plays an essential role in lipid transfer between these organelles (Guillen-Samander and De Camilli, 2022). VAP is present in two isoforms, VAPA and VAPB. VAPA is generally expressed in high levels compared to VAPB, so VAPA can compensate for the gene deletion of VAPB. So, we knocked down both VAPA and VAPB isoforms in our photoactivation and structural plasticity measurements. However, immunolocalization studies of the individual VAPA and VAPB isoforms reveal that VAPB is exclusively enriched near dendritic mitochondria, whereas VAPA is enriched near dendritic and axonal mitochondria. This result suggests that VAPB might be the dendrite-specific isoform specializing the tethering and stabilization rules of mitochondria in dendritic compartments. However, we did not find VAPB in our APEX-OMM proteome dataset, which might be due to its low expression levels compared to VAPA and its specific enrichment only near dendritic mitochondria. Furthermore, a P56S mutation in VAPB has been implicated in ALS (Nishimura et al., 2004). As most VAPB-related studies in ALS have focused only on axonal compartments, our novel findings on VAPB’s role in dendritic compartments might bring new perspectives to understanding ALS and the related deficiencies in synaptic plasticity and learning in motor neurons. Recently, VAPB has also been identified as a PD risk gene (Kun-Rodrigues et al., 2015). Future studies focusing on these mutations will illuminate their implications in synaptic plasticity in ALS and PD disease models.

ER-Mitochondria contact sites (ER-MCS) are crucial communication hotspots for ER-mitochondria calcium signaling, mitochondrial ATP production, and mitochondrial fission (Hirabayashi et al., 2017; Kozjak-Pavlovic, 2017; Simmen and Herrera-Cruz, 2018). Superresolution imaging of ER-MCS contacts in non-neuronal cells shows that nutrient deprivation reduces VAPB mobility at ER-MCS, potentially facilitating efficient trans-organelle metabolite transfer for signaling and energy production (Obara et al., 2022). We could therefore hypothesize that in neuronal dendrites at the base of the plasticity-induced spine, where energy demands are high, reduced VAP mobility might stabilize ER-actin-mitochondria interactions for long periods for enhanced, local energy production. Future studies investigating VAP dynamics and local energy production at ER-MCS in neuronal dendrites during synaptic plasticity will shed light on these questions.

Mitochondria represent a hugely underexplored organellar system in neurons. Notably, in dendritic compartments and during synaptic plasticity, the role of mitochondria has been largely understudied. Using recent advances in proteomics and imaging, we have discovered a novel function of VAP in spatially stabilizing mitochondria and its importance in fueling plasticity formation in stimulated and adjacent spines. These findings raise a new set of questions on the spatiotemporal regulation of local energy supplies and how they adapt to the local energetic needs of a dendritic segment. Understanding these specialized mechanisms also adds new perspectives to the compartment-specific rules of dendrites that are important for dendritic computation, learning, and memory formation (Harnett et al., 2012; Larkum and Nevian, 2008; Magee, 2000). Revealing these novel mitochondrial mechanisms will also bring new insights into therapeutic interventions for neurodegenerative disorders associated with mitochondrial dysfunction.

## Acknowledgments

We thank A.Y. Ting for the APEX2 plasmids, U. Manor for the Fis1-Lifeact-GFP and Fis1-mCherry plasmids, T. Meyer for the Ftractin-mCherry plasmid; E. Burton for the SNCA KD plasmid, F. Polleux for the SRGAP2 KD plasmid; C. Hoogenraad and M. Hoppa for the VAP KD and KO plasmids respectively; R. Youle for Mito-PAGFP plasmid; and J. Lippincott -Schwartz for VAPB-emerald plasmid. We are also thankful to E.M. Schuman for her support in the initial stages of this project; I. Wüllenweber for proteomics and mass spectrometry support; S. Perez, I. Bartnik, N. Fuerst, and A. Staab for technical assistance; Urban N for assistance with the MPFI Light Microscopy Core; and all members of the Rangaraju Lab, R. Yasuda, and H. Inagaki for critical input on the manuscript. VR is funded by the Max Planck Society, and TP was funded by the Nambu Summer Research Undergraduate Scholarship. This project has also received funding from the European Union’s Horizon 2020 research and innovation program under the Marie Sklodowska-Curie grant agreement 657702 for VR and the European Research Council grant agreement 743216 for E.M. Schuman.

## Author Contributions

VR designed experiments. Experiments were carried out by OB, TP, SK, CT, FR, and VR. Data were analyzed by OB, TP, SK, FR, MS, and VR. VR wrote the manuscript, and all authors edited the manuscript.

## Materials and Methods

### Animals

All experiments were performed according to the IACUC regulations.

### Plasmid constructs

APEX-Matrix (APEX2 targeted to peptide sequence derived from COX4) and APEX-OMM (APEX2 targeted to MAVS peptide sequence) were obtained from the Ting AY lab (Lam et al., 2015; Rhee et al., 2013). DsRed-Mito (Clontech) was purchased, and then DsRed was replaced with EGFP to make the EGFP-Mito construct. The Fis1-Lifeact-GFP and Fis1-mCherry plasmids were obtained from Manor U (Schiavon et al., 2020), SRGAP2 KD plasmids from Polleux F (Charrier et al., 2012), VAP KD and control plasmids from Hoogenraad C (Teuling et al., 2007), VAP KO and control sgRNA plasmids from Hoppa M (Panzera et al., 2022), and VAPB-emerald plasmid from Lippincott-Schwartz J. Ftractin-mCherry plasmid (Hayer et al., 2016), SNCA KD and control plasmids (Zharikov et al., 2015), and Mito-PAGFP plasmid (Karbowski et al., 2004) were purchased from Addgene (plasmids 85131, 75437, 75438, 23348 respectively). All other shRNAs against CAP1, IMMT, CYFIP1, NCKAP1, and PFN2 were purchased from Abbexa.

### Cell preparation, Transfection, Electroporation

Unless specified otherwise, all reagents were purchased from Sigma, and all stock solutions were stored at −20 °C. Sprague-Dawley rats (postnatal P0) were obtained from the in-house animal core facility of the Max Planck Florida Institute for Neuroscience. Hippocampal regions were dissected and dissociated with the Worthington Papain Dissociation Kit (Worthington Biochemical Corporation, stored at 4 °C) with a modified manufacturer’s protocol. Briefly, the digestion is done once for 30 min. Following trituration, the cells are centrifuged at 0.3 rcf for 5 minutes, resuspended in 2.4 ml of resuspension buffer (1.44 ml EBSS, 160 μl inhibitor, and 80 μl DNase) and 3.2 ml gradient, and then centrifuged at 0.6 rpm for 12 minutes. Neurons were plated on poly-D-lysine coated coverslips mounted on Mattek dishes at a density of 60,000-80,000 cells/cm^2^. Cultures were maintained in NGM (500 ml Neurobasal-A Medium, 10 ml B-27, and 5 ml Glutamax, from Gibco, ThermoFisher Scientific, sterile filtered using a 0.1 μm filter and stored at 4 °C) at 37 °C and 5% CO_2_. Cell feeding is done as follows: 3-4 hours after the cells are plated, they are fed with 1000 μl of NGM. Three days after, the cells are fed with 500 μl of NGM. Four days later, 300 μl of the media is removed from the dish and replenished with fresh 500 μl of NGM. Then, every 3-4 days, 300 μl is removed, and 500 μl is added until they are transfected. Transfections were performed 12-15 days after plating by magnetofection using Combimag (OZ biosciences) and Lipofectamine 2000 (Life Technologies) according to the manufacturer’s instructions.

For electroporation, the dissociated hippocampal cell suspension was obtained, as explained above, and neurons were electroporated using the Amaxa P3 primary cell 4D Nucleofector kit (Lonza) according to the manufacturer’s instructions. Each electroporation reaction (per electroporation cuvette) constituted 750,000 cells and 2 μg of APEX2-OMM DNA. 250,000 non-electroporated cells were later added to the electroporated cells to give a total of 1,000,000 cells and were plated on poly-D-lysine coated 6-cm plastic dishes (Greiner). Cultures were maintained, as explained above.

### APEX biotin labeling

Neurons co-transfected with either APEX-Matrix or APEX-OMM, along with EGFP-Mito, as a mitochondrial marker, were used for these experiments between 13-15 DIV. For biotin labeling, neurons were replaced with either 500 μM biotin phenol (Berry and Associates, 500 mM stock made in DMSO) or 500 μM biotin (500 mM stock made in DMSO stored at 4°C) in NGM for 30 min at 37 °C. Neurons were treated with 1 mM H_2_O_2_ (Alfa Aesar, 100 mM stock made in water and stored at 4 °C) for 1 min. The reaction was quenched by quencher solution consisting of (in mM) 10 Trolox (Merck), 20 sodium ascorbate, and 20 sodium azide (Roth) in DPBS, pH 7.4 adjusted with NaOH (filtered and stored at 4 °C), Neurons were subsequently washed thrice in either quencher wash solution consisting of (in mM) 5 Trolox, 10 sodium ascorbate, 10 sodium azide in DPBS for immunocytochemistry analyses or quencher solution for isolation of the biotinylated proteome.

### Immunostaining

Neurons were fixed in PFA (4% PFA in PBS) for 10 – 15 min, permeabilized (0.5% TritonX-100 in PBS for 30 min), and blocked in blocking buffer (4% goat serum in PBS for 1 hour). For VAP immunostaining alone, neurons were fixed in PFA (4% PFA in Cytoskeleton Buffer (CB) – in mM, 10 MES, pH 6.1, 138 KCl, 3 MgCl2, 2 EGTA, 0.3 M sucrose (Morley et al., 2016)) for 15 min, permeabilized (0.25% TritonX-100 in PBS for 15 min), and blocked in blocking buffer (2% BSA in PBS for 1 hour). Fixed and permeabilized neurons were incubated with anti-biotin rb (Bethyl, A150109A, 1:5000) for biotin labeling detection, anti-GFP chk (AVES 1020, 1:2000) for EGFP-Mito immunoamplification, anti-V5 ms (Invitrogen 460705, 1:500) for APEX2-Matrix immunoamplification, anti-FLAG ms (Sigma F3165, 1:500) for APEX2-OMM immunoamplification, anti-VAP A/B ms (NeuroMab 75-496, 1:100) or anti-VAPA rb (Proteintech 152751AP, 1:100), anti-VAPB rb (Proteintech 144771AP, 1:100) for VAP immunoamplification, and anti-SNCA ms (BD biosciences 610787, 1:1000) for SNCA immunoamplification for 90 min. Neurons were washed in PBS, and fluorophore-coupled secondary antibodies anti-rb Alexa 405 (Invitrogen A31556, 1:1000), anti chk Alexa 488 (Invitrogen A11039, 1:1000), anti-ms Alexa 546 (Invitrogen A11030, 1:1000), anti-ms Alexa 647 (Invitrogen A21236, 1:1000), and anti-rb Alexa 488 (Invitrogen A11008, 1:1000) were incubated for 45 – 60 min to detect biotin labeling, EGFP, APEX, VAP/SNCA, and VAPA/B respectively. Immunostained neurons were either stored at 4 °C or imaged directly.

### Biotinylated proteome isolation

For biotinylated proteome isolation, neurons were electroporated with APEX-OMM to achieve higher transfection efficiency. Following biotin labeling on 14 DIV neurons, as explained above, cells were scraped. For each experimental batch of neuronal culture preparation expressing APEX-OMM, cells from up to three 6 cm plastic dishes (adding up to 3,000,000 cells) were pooled together (to achieve ~70-100 μg of total protein). Cells were centrifuged at 600 rpm for 10 minutes at 4 °C, and the pellet was either stored at −80 °C or continued with cell lysis. Cell pellets were suspended in RIPA lysis buffer (RLB) consisting of (in mM) 50 Tris, 150 NaCl, 1 PMSF, 10 sodium azide, 10 sodium ascorbate, 5 Trolox, 0.01% SDS, 0.5% sodium deoxycholate, 1% Triton X-100 and 1X protease inhibitor cocktail in water, pH 7.4 (filtered and stored in 4 °C) and incubated in ice for 20 min. Lysed cells were centrifuged at 13000 rpm for 10 min at 4 °C, and the protein supernatant was collected. Protein concentration was determined using Pierce 660 nm protein assay (Thermofisher) according to the manufacturer’s instructions. Protein samples were stored at −80 °C.

### Streptavidin enrichment of biotinylated proteome

Biotinylated proteins were affinity-purified using streptavidin-coated magnetic beads (Pierce) as follows: Beads were washed twice in RLB, and protein samples were incubated with the beads (at 18:1 bead-to-sample ratio) for 1 hour at RT with gentle rotation (the flow through obtained here was used as FT). Beads were washed twice in RLB (the first wash was used as Wash), once in 2 M urea in 10 mM Tris-HCl, pH 8, and twice again in RLB. Beads were then washed thrice in RIPA wash buffer (RWB) containing (in mM) 50 Tris, 150 NaCl, 10 sodium azide, 10 sodium ascorbate, 5 Trolox, 0.1% SDS, 0.2% sodium deoxycholate and once with RWB2 containing (in mM) 50 Tris, 150 NaCl, 10 sodium azide, 10 sodium ascorbate, 5 Trolox and further processed for mass spectrometric analyses.

For western blot analyses, biotinylated proteins were eluted from the streptavidin beads using 3X NuPAGE LDS sample buffer supplemented with 20 mM DTT and 2 mM biotin and heating the mixture at 95 °C for 5 min. The elution step was performed twice to obtain Elu1 and Elu2 samples. Protein samples from input, FT, Wash, Elu1, and Elu2 were separated by electrophoresis (NuPAGE 4-12% Bis-Tris gel (Invitrogen)) and immunoblotted with anti-biotin ms (Sigma B7653, 1:2000) for detection of biotin labeling or stained with Coomassie for detection of total protein. Fluorophore-coupled secondary antibody anti-ms IR 800 (Invitrogen 92632210 LICOR, 1:10000) was used for immunodetection.

### Mass spectrometric analysis

Streptavidin-bound biotinylated proteins were digested on-bead using an optimized protocol (Schanzenbacher et al., 2016; Wisniewski et al., 2009). In brief, beads were washed 4 times in 1X PBS and twice in 50 mM ammonium bicarbonate, resuspended in 50 mM ammonium bicarbonate, and heated at 70 °C for 2 min with gentle agitation. 3 M urea was added, and the mixture was cooled down to RT. Bead suspension was reduced with 3.125 mM TCEP (0.5 M stock prepared in LC-MS-grade water) for 30 min at RT with constant agitation and incubated with 11.2 mM iodoacetamide (0.5 M stock prepared in LC-MS-grade water) for 30 min at RT in the dark with continuous agitation. Protein samples were digested sequentially with Lys-C and trypsin on Microcon-10 filters (Merck Millipore, #MRCPRT010 Ultracel YM-10). The mixture was centrifuged at 1000 rcf for 5 min, and the supernatant containing peptides was collected. Digested peptides were desalted using ZipTips (Merck) according to the manufacturer’s instructions, dried in a Speed-Vac, and stored at −20 °C. For mass spectrometry analyses, the dried peptides were reconstituted in 5% acetonitrile with 0.1% formic acid and loaded using a nano-HPLC (Dionex U3000 RSLCnano) on reversed-phase columns (trapping column: particle size = 3 μm, C18, L = 2 cm; analytical column: particle size = 2 μm, C18, L = 50 cm; PepMap, Dionex/Thermofisher). Peptides eluting from the column were ionized using an Orbitrap Elite (Thermofisher).

First, a full MS scan from m/z 350 to1600 was acquired at a resolution of 120,000, and sequence information was acquired by computer-controlled, data-dependent automated switching to MS/MS mode using collision-induced dissociation (CID) fragmentation with normal collision energy (NCE) of 35 % in the linear Ion Trap (IT-MS). Next, MSMS-spectra were searched against the protein database for Rattus norvegicus (TaxID 10116, Uniprot) and additionally for a database containing common MS-contaminations with trypsin as digestion enzyme using the Proteome Discoverer (PD) (Thermo Fisher, version 2.1). The set of stringent search constraints allowed tryptic peptides with up to two missed cleavages, a minimum of two valid peptides per locus, a precursor mass tolerance of 10 ppm, and a fragment mass tolerance of 0.5 Da. In addition, carbamidomethylation of cysteine was set as a fixed modification, acetylation of peptide N-termini, oxidation of methionine, and biotin-labeling of tyrosine, tryptophan, and histidine were set as variable modifications. Percolator node, a machine-learning tool in PD 2.1, was used to estimate the number of false positive identifications. A high confidence q-value of less than 0.01 (False discovery rate (FDR) < 0.01) was assigned to filter both the peptide spectrum match (PSM) results and peptide results.

The procedure was repeated thrice, and three technical replicates were combined. The technical overlap for high confident proteins was determined, where at least two peptides from Rattus norvegicus were needed for identification. Proteins found in the control sample were excluded from the experimental samples. Proteins found in at least two of the three replicates were considered for further analysis. The Gene/product IDs for these proteins were searched through Gene Ontology Resource for ‘mitochondrion’ as GO cellular component with *Rattus norvegicus* as a filter.

The Gene/product IDs were further filtered for actin and tubulin interactors using a database for protein interactions, BioGRID, using the search term ‘Actb’ for actin interactors and ‘Tuba1a’ for tubulin interactors with *Rattus norvegicus* and *Mus musculus* as organism filters.

The mass spectrometry proteomics data have been deposited to the ProteomeXchange Consortium (http://proteomecentral.proteomexchange.org) via the PRIDE partner repository (Perez-Riverol et al., 2019). Anonymous reviewer access is available upon request.

### Imaging

Live cell imaging was conducted 15-18 days after neuronal cell culture plating. All experiments, unless specified, were performed at 32 °C in E4 imaging buffer containing (in mM) 120 NaCl, 3 KCl, 10 HEPES (buffered to pH 7.4), 3 CaCl_2_, 1 MgCl_2_, and 10 Glucose. For spine stimulation by glutamate uncaging experiments, a modified E4 buffer lacking MgCl_2_ with 4 mM CaCl_2_ was used –the rest of the constituents remained unchanged. Airyscan confocal experiments were done at room temperature.

### Optical Measurements

Live imaging was performed using a custom-built inverted spinning disk confocal microscope (3i imaging systems; model CSU-W1) attached to an Andor iXon Life 888. Image acquisition was controlled by SlideBook 6 software. Images were acquired with a Plan-Apochromat 63x/1.4 NA. Oil objective, M27 with DIC III prism, using a CSU-W1 Dichroic for 488/561 nm excitation with Quad emitter and individual emitters, at laser powers 1.1 mW (488 nm) and 0.8 mW (561 nm) for photoactivation and VAPB OE (VAPB-emerald) experiments; and laser powers 2 mW (488 nm) and 2.7 mW (561 nm) for spine structural plasticity measurements. During imaging, the temperature was maintained at 32 °C using an Okolab stage top incubator with temperature control.

### Mitochondria-Actin Interaction Measurements using Airyscan Confocal

Neurons were transfected with Fis1-Lifeact-GFP and Fis1-mCherry plasmid constructs, along with either control shRNA or shRNA targeting a gene of interest (Cap1, Immt, Snca, Cyfip1, Nckap1, Pfn2, Srgap2, Vap, or Mcu). 3-7 days post-transfection, neurons were identified by Fis1-mCherry signal targeting mitochondria to differentiate between dendritic (long mitochondria) and axonal (short mitochondria) regions. For negative control experiments, neurons transfected with Fis1-Lifeact-GFP and Fis1-mCherry plasmid constructs were imaged either in the absence or presence of 25 μg/ml cytochalasin-D for 15 min (12.5 mg/ml stock made in DMSO), 1 μg/ml Latrunculin-B for 10 min (1 mg/ml stock made in DMSO), or 10 μg/ml nocodazole for 10 min (5 mg/ml stock made in DMSO). Neurons were transfected with Ftractin-mCherry in the absence or presence of Fis1-Lifeact-GFP to visualize any effect of Lifeact-GFP expression on actin morphology in dendrites.

Imaging was performed using an LSM 880 AxioObserver confocal microscope (Zeiss) with a Plan-Apochromat 63x/1.4 DIC oil immersion objective (Zeiss) and the Airyscan (GaAsP-PMT) detector. Images were acquired in 16-bit mode using unidirectional scanning, with a field of view of 66.45 μm x 66.45 μm (zoom 2.0), voxel sizes of (x,y,z) 40 nm x 40 nm x 800 nm, and pixel dwell times of 2.65 μs. The height (along the optical axis) of the volumetric z-stack was chosen to encompass the entire thickness of the neuron. Dual-color Airyscan imaging was performed sequentially using frame steps in Airyscan-SR mode with a default pinhole size set to 2.77 AU (for 488 nm excitation) and 2.5 AU (for 561 nm excitation). Laser power and detector gain were adjusted for each channel to utilize between 35% – 50% of the full dynamic range of the detector, carefully avoiding detector saturation. Typical laser powers were around 5.0 % (488 nm) and 4.5 % (561 nm) with a master gain set to 750 V. Raw Airyscan images were subsequently processed using Zeiss Airyscan processing set to default deconvolution strength settings. Imaging conditions were kept constant within a batch of experiments.

The Imaris software package (version 9.5.1; Oxford Instruments/Andor) was used to analyze the processed Airyscan images of the mitochondria-actin interaction experiments. First, two different surfaces were created, one for the red color channel (Fis1-mCherry) and one for the green color channel (Fis1-Lifeact-GFP). Then, a colocalized surface was made in yellow using the ‘colocalization’ feature. Next, total volume was obtained from the red channel surface (total mitochondria) and the yellow colocalized surface (mitochondrial region interacting with actin) using the ‘detailed statistics’ feature. Finally, the percentage interaction between mitochondria and actin was calculated by dividing the volume of the yellow colocalized surface (mitochondrial region interacting with actin) by the volume of the red channel surface (total mitochondria).

### Photoactivation experiments

A custom-written automated program was used for simultaneous imaging of up to 10 neurons, per dish, with a z-stack interval of 1 μm spanning a total of 5 μm, with both 488 nm and 561 nm lasers at exposure times of 15 ms. Neurons were transfected with Mito-PAGFP and Mito-DsRed plasmid constructs along with control shRNA or shRNA plasmids targeting the desired gene (Vapa/b, Snca, or Srgap2). Transfected neurons were identified using Mito-DsRed fluorescence. Photoactivation was performed using a multi-photon laser set-up at 805 nm (Chameleon, Coherent) and a Pockels cell (Conoptics) for controlling the photoactivation pulse. A 5×2 – 10×2 μm^2^ photoactivation spot was chosen at a dendrite. Images were acquired every 30 s for 70 min. Following the acquisition of pre-photoactivation fluorescence images for the first 6 time frames (3 min), a photoactivation pulse was given at the 7th time frame at 1 ms duration per pixel, and 3.5 mW power and the imaging continued.

For measuring the photoactivated mitochondrial compartment length from these experiments, images were converted to maximum intensity projections and analyzed by Imaris 8.4.1. Using the ‘surface segmentation wizard’, the photoactivated mitochondrial region was selected by setting a manual threshold, excluding the non-photoactivated background. The ‘compartment length’ was defined as the photoactivated mitochondria extracted across all the time points using the ‘filament wizard’ and the subsequent addition of the photoactivated mitochondrial filaments detected above the set threshold during the imaging period.

For measuring the photoactivated mitochondrial compartment fluorescence from these experiments, an ROI of the size of the photoactivated mitochondrial compartment at the 5 min timepoint was drawn, and the fluorescence intensity was measured over the 60 min time course of the experiment. The photoactivated compartment stability index was calculated by dividing the photoactivated compartment fluorescence intensity at the end of 60 min postphotoactivation (F_60min_) by the photoactivated compartment fluorescence intensity 5 min postphotoactivation (F_5min_), resulting in F_60min_/F_5min_. A reduced compartment stability index statistically significant from the control was considered an unstable mitochondrial compartment.

### Correlation coefficient measurements

Neurons transfected with Mito-DsRed were fixed and immunostained for VAPA or VAPB, as explained above. Images were acquired with an LSM 880 confocal microscope (Zeiss) using a C-Apochromat 40x/1.2W objective and a pinhole size of 1 airy unit. Images were acquired in 16-bit mode as Z stacks, with a pixel pitch of 208 nm and field of view of 212.55 × 212.55 μm^2^, throughout the entire thickness of the neuron with an optical slice thickness of 0.5 μm and pixel dwell times of 0.5 μs. The detector gain in each channel was adjusted to cover the full dynamic range, avoiding saturated pixels. Imaging conditions were kept constant within a batch of experiments. Maximum intensity projections were used for image analyses.

For live imaging, neurons were transfected with Mito-DsRed and VAPB-emerald. Images were acquired from the dendritic and axonal compartments using the spinning disc confocal microscope, as explained above.

For correlation coefficient analysis, dendritic and axonal regions of interest from both immunostained and live neuron images were straightened from the respective VAPA/B (green) and mitochondria (red) channels using ImageJ. A line of ~3 μm thickness was drawn along the straightened dendrite or axon from the respective channels, and a line profile was obtained. Each dendrite or axon’s VAPA/B line profile was plotted against its mitochondria line profile, and the corresponding Pearson correlation coefficients were determined using OriginPro 2022 (64-bit) SR1. For shuffled control (Shf), one of the line profile measurements (mitochondria channel) was shuffled, while the corresponding line profile pair (VAP channel) remained undisturbed, and the corresponding Pearson correlation coefficient was determined.

### Spine structural plasticity measurements

Neurons were transfected with GCaMP6s, Mito-DsRed, and PSD95-mCherry or Homer2-mOrange plasmid constructs, along with Control Cas9 plasmid lacking sgRNAs or with the VAP CRISPR-Cas9 KO plasmid with two sgRNAs targeting Vapa and Vapb isoforms, respectively. Transfected neurons were identified by a change in GCaMP6s fluorescence corresponding to calcium transients in dendrites and spines. In addition, PSD95-mCherry or Homer2-mOrange fluorescence was used to identify spines for stimulation by two-photon glutamate uncaging or to identify adjacent spines in the same dendritic segment or sister dendrites. Before glutamate uncaging, neurons were replaced with 1 μM TTX (Citrate salt, 2 mM stock made in water), 50 μM Forskolin (Tocris Bioscience, 25 mM stock made in DMSO), and 2 mM 4-Methoxy-7-nitroindolinyl-caged-L-glutamate (MNI caged glutamate) (Tocris Bioscience, 100 mM stock made in E4 buffer) in modified E4 buffer lacking Mg^2+^ (see above). Glutamate uncaging was performed using a multiphoton-laser 720 nm (Mai TAI HP) and a Pockel cell (ConOptics) for controlling the uncaging pulses. Spines that were at least 50 μm away from the cell body were chosen for uncaging experiments. To test a spine’s response to an uncaging pulse, an uncaging spot (2 μm^2^) close to a spine-head was selected, and two to three uncaging pulses at 10 ms pulse duration per pixel and 1.2 mW power were given and checked for spine-specific calcium transients. An uncaging protocol of 60 uncaging pulses at 0.5 Hz with 10 ms pulse duration per pixel at 1.2 mW power was used. Images were acquired before plasticity induction at t = 0 min; at the end of the plasticity induction at t = 2 min; and then every 10 min for up to 62 min at t = 12, 22, 32, 42, 52, 62 min.

Following structural plasticity measurements, neurons were fixed in CB buffer as explained above, stored either for 3-4 hours at room temperature (or at 4 °C overnight), and continued with the VAP immunostaining protocol (see above).

Using a custom-written FIJI Macro, ten images from each timepoint were averaged, and the background was subtracted using a rolling ball filter of a radius 50 pixels. Next, a line crossing the center of the spine-head was drawn using a custom-written MATLAB script. Then, the fluorescence intensity measured along the line was fit to a Gaussian to obtain the full width at half maxima (FWHM) –defined as the spine-head width (Matsuzaki et al., 2004). As the FWHM is independent of fluorescence intensity, it was used to measure spine-head width even from fluctuating GCaMP fluorescence in spines.

### shRNA knockdown efficiency determination

Following photoactivation or spine structural plasticity measurements, the neurons were fixed and immunostained, as explained above. Expression levels were quantified by selecting soma regions of interest (ROIs) located by either GCaMP signal (from structural plasticity measurements) or Mito-PAGFP signal (from photoactivation experiments) followed by measuring VAP, SNCA, or any other protein’s immunofluorescence intensity at the same ROIs after background subtraction. The average VAP or SNCA fluorescence intensities measured in GCaMP-positive or Mito-PAGFP-positive, transfected somas (F_trans_) and the average VAP or SNCA fluorescence intensities measured in GCaMP-negative or Mito-PAGFP-negative somas from adjacent untransfected neurons (F_untrans_) were obtained from the same illumination field. The knockdown percentage was calculated as F_trans_/F_untrans_.

### Quantification and statistical analysis

Statistical details of experiments, including statistical tests and n and p values, can be found in their respective figure legends. In all the figures, the box whisker plots represent the median (line), mean (point), 25th-75th percentile (box), 10th-90th percentile (whisker), 1st–99th percentile (X), and min-max (_) ranges. Error bars in bar graphs and line traces are SEM. A p-value of less than or equal to 0.05 was considered significant for all statistical tests.

## References

Basu, H., Pekkurnaz, G., Falk, J., Wei, W., Chin, M., Steen, J., and Schwarz, T.L. (2021). FHL2 anchors mitochondria to actin and adapts mitochondrial dynamics to glucose supply. The Journal of cell biology 220.

Bradshaw, K.D., Emptage, N.J., and Bliss, T.V. (2003). A role for dendritic protein synthesis in hippocampal late LTP. Eur J Neurosci 18, 3150–3152.

Charrier, C., Joshi, K., Coutinho-Budd, J., Kim, J.E., Lambert, N., de Marchena, J., Jin, W.L., Vanderhaeghen, P., Ghosh, A., Sassa, T., et al. (2012). Inhibition of SRGAP2 function by its human-specific paralogs induces neoteny during spine maturation. Cell 149, 923–935.

Courchet, J., Lewis, T.L., Jr., Lee, S., Courchet, V., Liou, D.Y., Aizawa, S., and Polleux, F. (2013). Terminal axon branching is regulated by the LKB1-NUAK1 kinase pathway via presynaptic mitochondrial capture. Cell 153, 1510–1525.

Dong, R., Saheki, Y., Swarup, S., Lucast, L., Harper, J.W., and De Camilli, P. (2016). Endosome-ER Contacts Control Actin Nucleation and Retromer Function through VAP-Dependent Regulation of PI4P. Cell 166, 408–423.

Faits, M.C., Zhang, C.M., Soto, F., and Kerschensteiner, D. (2016). Dendritic mitochondria reach stable positions during circuit development. Elife 5.

Fu, M., Yu, X., Lu, J., and Zuo, Y. (2012). Repetitive motor learning induces coordinated formation of clustered dendritic spines in vivo. Nature 483, 92–95.

Gomez-Suaga, P., Perez-Nievas, B.G., Glennon, E.B., Lau, D.H.W., Paillusson, S., Morotz, G.M., Cali, T., Pizzo, P., Noble, W., and Miller, C.C.J. (2019). The VAPB-PTPIP51 endoplasmic reticulum-mitochondria tethering proteins are present in neuronal synapses and regulate synaptic activity. Acta Neuropathol Commun 7, 35.

Govindarajan, A., Israely, I., Huang, S.Y., and Tonegawa, S. (2011). The dendritic branch is the preferred integrative unit for protein synthesis-dependent LTP. Neuron 69, 132–146.

Govindarajan, A., Kelleher, R.J., and Tonegawa, S. (2006). A clustered plasticity model of long-term memory engrams. Nature reviews Neuroscience 7, 575–583.

Guillen-Samander, A., and De Camilli, P. (2022). Endoplasmic Reticulum Membrane Contact Sites, Lipid Transport, and Neurodegeneration. Cold Spring Harbor perspectives in biology.

Harnett, M.T., Makara, J.K., Spruston, N., Kath, W.L., and Magee, J.C. (2012). Synaptic amplification by dendritic spines enhances input cooperativity. Nature 491, 599–602.

Hayer, A., Shao, L., Chung, M., Joubert, L.M., Yang, H.W., Tsai, F.C., Bisaria, A., Betzig, E., and Meyer, T. (2016). Engulfed cadherin fingers are polarized junctional structures between collectively migrating endothelial cells. Nature cell biology 18, 1311–1323.

Hirabayashi, Y., Kwon, S.K., Paek, H., Pernice, W.M., Paul, M.A., Lee, J., Erfani, P., Raczkowski, A., Petrey, D.S., Pon, L.A., et al. (2017). ER-mitochondria tethering by PDZD8 regulates Ca(2+) dynamics in mammalian neurons. Science 358, 623–630.

Holt, C.E., Martin, K.C., and Schuman, E.M. (2019). Local translation in neurons: visualization and function. Nat Struct Mol Biol 26, 557–566.

Hubley, M.J., Locke, B.R., and Moerland, T.S. (1996). The effects of temperature, pH, and magnesium on the diffusion coefficient of ATP in solutions of physiological ionic strength. Biochimica et biophysica acta 1291, 115–121.

Hung, V., Lam, S.S., Udeshi, N.D., Svinkina, T., Guzman, G., Mootha, V.K., Carr, S.A., and Ting, A.Y. (2017). Proteomic mapping of cytosol-facing outer mitochondrial and ER membranes in living human cells by proximity biotinylation. Elife 6.

Jang, S., Nelson, J.C., Bend, E.G., Rodriguez-Laureano, L., Tueros, F.G., Cartagenova, L., Underwood, K., Jorgensen, E.M., and Colon-Ramos, D.A. (2016). Glycolytic Enzymes Localize to Synapses under Energy Stress to Support Synaptic Function. Neuron 90, 278–291.

Kang, H., and Schuman, E.M. (1996). A requirement for local protein synthesis in neurotrophin-induced hippocampal synaptic plasticity. Science 273, 1402–1406.

Kang, J.S., Tian, J.H., Pan, P.Y., Zald, P., Li, C., Deng, C., and Sheng, Z.H. (2008). Docking of axonal mitochondria by syntaphilin controls their mobility and affects short-term facilitation. Cell 132, 137–148.

Karbowski, M., Arnoult, D., Chen, H., Chan, D.C., Smith, C.L., and Youle, R.J. (2004). Quantitation of mitochondrial dynamics by photolabeling of individual organelles shows that mitochondrial fusion is blocked during the Bax activation phase of apoptosis. The Journal of cell biology 164, 493–499.

Kleindienst, T., Winnubst, J., Roth-Alpermann, C., Bonhoeffer, T., and Lohmann, C. (2011). Activity-dependent clustering of functional synaptic inputs on developing hippocampal dendrites. Neuron 72, 1012–1024.

Kozjak-Pavlovic, V. (2017). The MICOS complex of human mitochondria. Cell Tissue Res 367, 83–93.

Kun-Rodrigues, C., Ganos, C., Guerreiro, R., Schneider, S.A., Schulte, C., Lesage, S., Darwent, L., Holmans, P., Singleton, A., International Parkinson’s Disease Genomics, C., et al. (2015). A systematic screening to identify de novo mutations causing sporadic early-onset Parkinson’s disease. Hum Mol Genet 24, 6711–6720.

Lam, S.S., Martell, J.D., Kamer, K.J., Deerinck, T.J., Ellisman, M.H., Mootha, V.K., and Ting, A.Y. (2015). Directed evolution of APEX2 for electron microscopy and proximity labeling. Nature methods 12, 51–54.

Larkum, M.E., and Nevian, T. (2008). Synaptic clustering by dendritic signalling mechanisms. Curr Opin Neurobiol 18, 321–331.

Lee, A., Hirabayashi, Y., Kwon, S.K., Lewis, T.L., Jr., and Polleux, F. (2018). Emerging roles of mitochondria in synaptic transmission and neurodegeneration. Curr Opin Physiol 3, 82–93.

Lewis, T.L., Jr., Kwon, S.K., Lee, A., Shaw, R., and Polleux, F. (2018). MFF-dependent mitochondrial fission regulates presynaptic release and axon branching by limiting axonal mitochondria size. Nat Commun 9, 5008.

Lewis, T.L., Jr., Turi, G.F., Kwon, S.K., Losonczy, A., and Polleux, F. (2016). Progressive Decrease of Mitochondrial Motility during Maturation of Cortical Axons In Vitro and In Vivo. Curr Biol 26, 2602–2608.

Li, Z., Okamoto, K., Hayashi, Y., and Sheng, M. (2004). The importance of dendritic mitochondria in the morphogenesis and plasticity of spines and synapses. Cell 119, 873–887.

Ligon, L.A., and Steward, O. (2000). Role of microtubules and actin filaments in the movement of mitochondria in the axons and dendrites of cultured hippocampal neurons. The Journal of comparative neurology 427, 351–361.

Lindhout, F.W., Cao, Y., Kevenaar, J.T., Bodzeta, A., Stucchi, R., Boumpoutsari, M.M., Katrukha, E.A., Altelaar, M., MacGillavry, H.D., and Hoogenraad, C.C. (2019). VAP-SCRN1 interaction regulates dynamic endoplasmic reticulum remodeling and presynaptic function. The EMBO journal 38, e101345.

MacAskill, A.F., Rinholm, J.E., Twelvetrees, A.E., Arancibia-Carcamo, I.L., Muir, J., Fransson, A., Aspenstrom, P., Attwell, D., and Kittler, J.T. (2009). Miro1 Is a Calcium Sensor for Glutamate Receptor-Dependent Localization of Mitochondria at Synapses. Neuron 61, 541–555.

Magee, J.C. (2000). Dendritic integration of excitatory synaptic input. Nature reviews Neuroscience 1, 181–190.

Makino, H., and Malinow, R. (2011). Compartmentalized versus global synaptic plasticity on dendrites controlled by experience. Neuron 72, 1001–1011.

Martin, K.C., Casadio, A., Zhu, H., Yaping, E., Rose, J.C., Chen, M., Bailey, C.H., and Kandel, E.R. (1997). Synapse-specific, long-term facilitation of aplysia sensory to motor synapses: a function for local protein synthesis in memory storage. Cell 91, 927–938.

Matsuzaki, M., Honkura, N., Ellis-Davies, G.C., and Kasai, H. (2004). Structural basis of long-term potentiation in single dendritic spines. Nature 429, 761–766.

McBride, T.J., Rodriguez-Contreras, A., Trinh, A., Bailey, R., and Debello, W.M. (2008). Learning drives differential clustering of axodendritic contacts in the barn owl auditory system. The Journal of neuroscience: the official journal of the Society for Neuroscience 28, 6960–6973.

Morley, S.J., Qi, Y., Iovino, L., Andolfi, L., Guo, D., Kalebic, N., Castaldi, L., Tischer, C., Portulano, C., Bolasco, G., et al. (2016). Acetylated tubulin is essential for touch sensation in mice. Elife 5.

Nishimura, A.L., Mitne-Neto, M., Silva, H.C., Richieri-Costa, A., Middleton, S., Cascio, D., Kok, F., Oliveira, J.R., Gillingwater, T., Webb, J., et al. (2004). A mutation in the vesicle-trafficking protein VAPB causes late-onset spinal muscular atrophy and amyotrophic lateral sclerosis. Am J Hum Genet 75, 822–831.

Nishiyama, J., and Yasuda, R. (2015). Biochemical Computation for Spine Structural Plasticity. Neuron 87, 63–75.

Obara, C.J., Nixon-Abell, J., Moore, A.S., Riccio, F., Hoffman, D.P., Shtengel, G., Xu, C.S., Schaefer, K., Pasolli, H.A., Masson, J.-B., et al. (2022). Motion of single molecular tethers reveals dynamic subdomains at ER-mitochondria contact sites. bioRxiv, 2022.2009.2003.505525.

Oughtred, R., Rust, J., Chang, C., Breitkreutz, B.J., Stark, C., Willems, A., Boucher, L., Leung, G., Kolas, N., Zhang, F., et al. (2021). The BioGRID database: A comprehensive biomedical resource of curated protein, genetic, and chemical interactions. Protein Sci 30, 187–200.

Panzera, L.C., Johnson, B., Quinn, J.A., Cho, I.H., Tamkun, M.M., and Hoppa, M.B. (2022). Activity-dependent endoplasmic reticulum Ca(2+) uptake depends on Kv2.1-mediated endoplasmic reticulum/plasma membrane junctions to promote synaptic transmission. Proceedings of the National Academy of Sciences of the United States of America 119, e2117135119.

Pathak, D., Shields, L.Y., Mendelsohn, B.A., Haddad, D., Lin, W., Gerencser, A.A., Kim, H., Brand, M.D., Edwards, R.H., and Nakamura, K. (2015). The role of mitochondrially derived ATP in synaptic vesicle recycling. J Biol Chem 290, 22325–22336.

Perez-Riverol, Y., Csordas, A., Bai, J., Bernal-Llinares, M., Hewapathirana, S., Kundu, D.J., Inuganti, A., Griss, J., Mayer, G., Eisenacher, M., et al. (2019). The PRIDE database and related tools and resources in 2019: improving support for quantification data. Nucleic Acids Res 47, D442–D450.

Popov, V., Medvedev, N.I., Davies, H.A., and Stewart, M.G. (2005). Mitochondria form a filamentous reticular network in hippocampal dendrites but are present as discrete bodies in axons: a three-dimensional ultrastructural study. The Journal of comparative neurology 492, 50–65.

Rangaraju, V., Calloway, N., and Ryan, T.A. (2014). Activity-driven local ATP synthesis is required for synaptic function. Cell 156, 825–835.

Rangaraju, V., Lauterbach, M., and Schuman, E.M. (2019a). Spatially Stable Mitochondrial Compartments Fuel Local Translation during Plasticity. Cell 176, 73–84 e15.

Rangaraju, V., Lewis, T.L., Jr., Hirabayashi, Y., Bergami, M., Motori, E., Cartoni, R., Kwon, S.K., and Courchet, J. (2019b). Pleiotropic Mitochondria: The Influence of Mitochondria on Neuronal Development and Disease. The Journal of neuroscience: the official journal of the Society for Neuroscience 39, 8200–8208.

Rangaraju, V., Tom Dieck, S., and Schuman, E.M. (2017). Local translation in neuronal compartments: how local is local? EMBO Rep 18, 693–711.

Rhee, H.W., Zou, P., Udeshi, N.D., Martell, J.D., Mootha, V.K., Carr, S.A., and Ting, A.Y. (2013). Proteomic mapping of mitochondria in living cells via spatially restricted enzymatic tagging. Science 339, 1328–1331.

Riedl, J., Crevenna, A.H., Kessenbrock, K., Yu, J.H., Neukirchen, D., Bista, M., Bradke, F., Jenne, D., Holak, T.A., Werb, Z., et al. (2008). Lifeact: a versatile marker to visualize F-actin. Nature methods 5, 605–607.

Schanzenbacher, C.T., Sambandan, S., Langer, J.D., and Schuman, E.M. (2016). Nascent Proteome Remodeling following Homeostatic Scaling at Hippocampal Synapses. Neuron 92, 358–371.

Schiavon, C.R., Zhang, T., Zhao, B., Moore, A.S., Wales, P., Andrade, L.R., Wu, M., Sung, T.C., Dayn, Y., Feng, J.W., et al. (2020). Actin chromobody imaging reveals sub-organellar actin dynamics. Nature methods 17, 917–921.

Shepherd, G.M., and Harris, K.M. (1998). Three-dimensional structure and composition of CA3-->CA1 axons in rat hippocampal slices: implications for presynaptic connectivity and compartmentalization. The Journal of neuroscience: the official journal of the Society for Neuroscience 18, 8300–8310.

Simmen, T., and Herrera-Cruz, M.S. (2018). Plastic mitochondria-endoplasmic reticulum (ER) contacts use chaperones and tethers to mould their structure and signaling. Curr Opin Cell Biol 53, 61–69.

Smith, H.L., Bourne, J.N., Cao, G., Chirillo, M.A., Ostroff, L.E., Watson, D.J., and Harris, K.M. (2016). Mitochondrial support of persistent presynaptic vesicle mobilization with age-dependent synaptic growth after LTP. Elife 5.

Spillane, M., Ketschek, A., Merianda, T.T., Twiss, J.L., and Gallo, G. (2013). Mitochondria coordinate sites of axon branching through localized intra-axonal protein synthesis. Cell Rep 5, 1564–1575.

Takahashi, N., Kitamura, K., Matsuo, N., Mayford, M., Kano, M., Matsuki, N., and Ikegaya, Y. (2012). Locally synchronized synaptic inputs. Science 335, 353–356.

Teuling, E., Ahmed, S., Haasdijk, E., Demmers, J., Steinmetz, M.O., Akhmanova, A., Jaarsma, D., and Hoogenraad, C.C. (2007). Motor neuron disease-associated mutant vesicle-associated membrane protein-associated protein (VAP) B recruits wild-type VAPs into endoplasmic reticulum-derived tubular aggregates. The Journal of neuroscience: the official journal of the Society for Neuroscience 27, 9801–9815.

Vickers, C.A., Dickson, K.S., and Wyllie, D.J. (2005). Induction and maintenance of late-phase long-term potentiation in isolated dendrites of rat hippocampal CA1 pyramidal neurones. J Physiol 568, 803–813.

Wang, X.N., and Schwarz, T.L. (2009). The Mechanism of Ca2+-Dependent Regulation of Kinesin-Mediated Mitochondrial Motility. Cell 136, 163–174.

Wang, Y., Xu, E., Musich, P.R., and Lin, F. (2019). Mitochondrial dysfunction in neurodegenerative diseases and the potential countermeasure. CNS Neurosci Ther 25, 816–824.

Wilson, D.E., Whitney, D.E., Scholl, B., and Fitzpatrick, D. (2016). Orientation selectivity and the functional clustering of synaptic inputs in primary visual cortex. Nat Neurosci 19, 1003–1009.

Wisniewski, J.R., Zougman, A., Nagaraj, N., and Mann, M. (2009). Universal sample preparation method for proteome analysis. Nature methods 6, 359–362.

Zharikov, A.D., Cannon, J.R., Tapias, V., Bai, Q., Horowitz, M.P., Shah, V., El Ayadi, A., Hastings, T.G., Greenamyre, J.T., and Burton, E.A. (2015). shRNA targeting alpha-synuclein prevents neurodegeneration in a Parkinson’s disease model. J Clin Invest 125, 2721–2735.

