## Supplementary Figures for "VAP spatially stabilizes dendritic mitochondria to locally fuel synaptic plasticity"

### Supplementary Figures and Legends

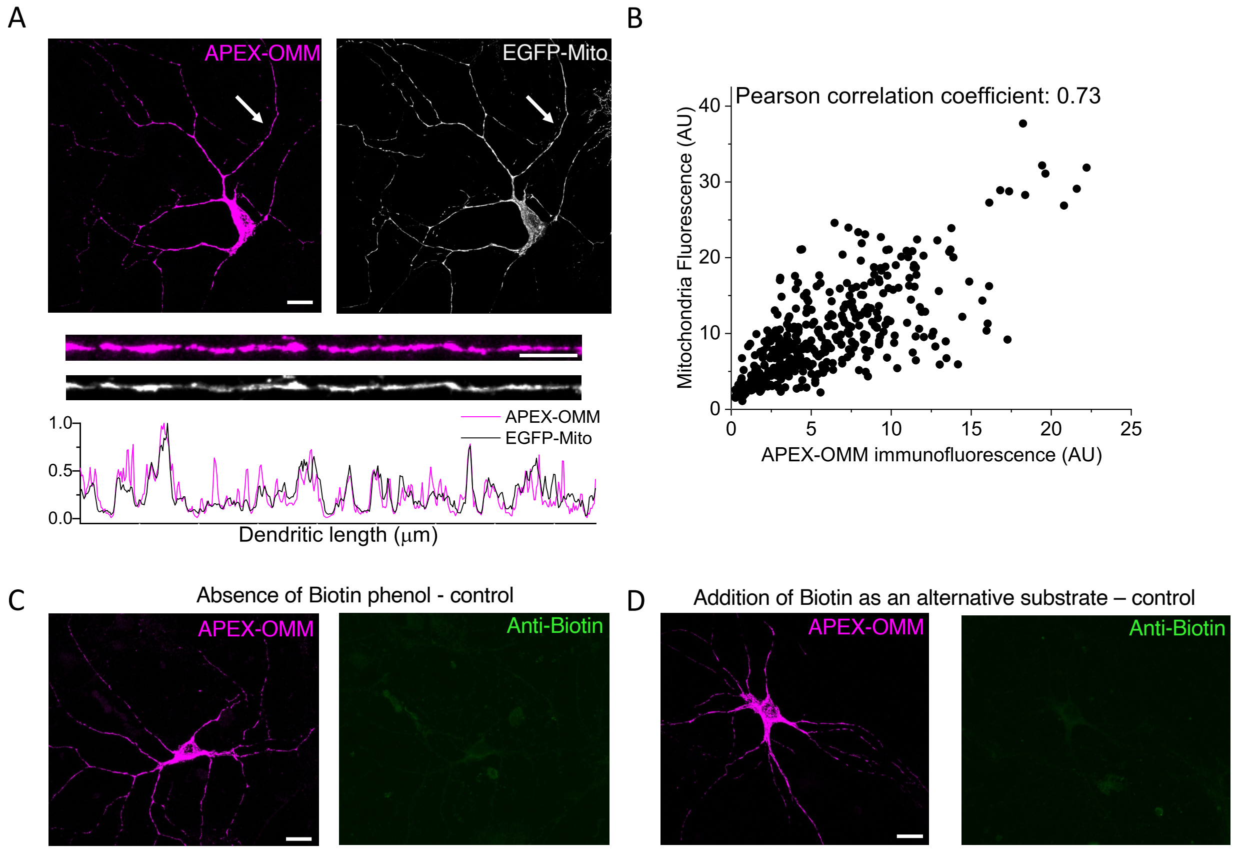


**Figure S1. Related to Figure 1. Control experiments related to APEX-OMM expression and biotin labeling**

A. (Top) APEX-OMM fluorescence (left, same as in Figure 1B) and mitochondria fluorescence (right, EGFP-Mito) from the same neuron overlap. Scale bar, 20 μm. (Bottom) Line profiles of the dendrites pointed in A (white arrows) show an overlap between APEX-OMM and mitochondria fluorescence. B. Representative correlation between individual fluorescent pixel intensities of APEX-OMM and mitochondria in Fig. S1A shows a strong correlation. Representative neuron images expressing APEX-OMM (left, magenta), showing the absence of biotin labeling (right, green) in the absence of biotin phenol (C) or the presence of biotin as an alternative substrate (D). Scale bar, 10 μm.

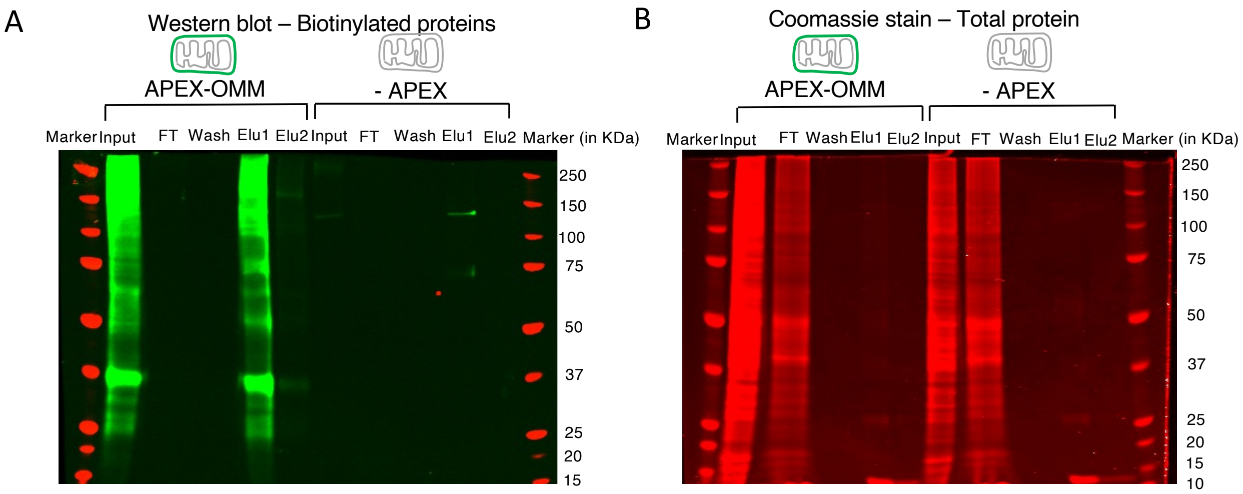


**Figure S2. Related to Figure 2. Streptavidin enrichment of the biotin-labeled APEX-OMM proteome**

(A) Western blot analysis of the biotin-labeled, streptavidin-enriched proteome using anti-biotin immunodetection (see Methods) showed significant biotin labeling in the input (Input) and eluate 1 (Elu1) of APEX-OMM samples but not in -APEX (control) samples. The residual biotin signal in control represents endogenous biotinylated proteins present in mammalian cells (Chapman-Smith and Cronan, 1999). (B) Coomassie staining analysis of the same experiment showed equal amounts of protein input used for APEX-OMM and -APEX (control)samples, and most non-biotinylated proteins were washed out in the flow through (FT, see Methods).

**Table S1. Related to Figure 2. The neuronal OMM proteome comprises 129 proteins.**

Column 1: UniProt ID of the OMM protome

Column 2: Gene names of the OMM proteome

Column 3: Protein names of the OMM proteome

Columns 4, 5, 6: Presence or absence of the OMM proteome in replicates 1, 2, and 3, respectively.

Column 7: GO annotation of the OMM proteome with the term “Mitochondria” reveals 26 proteins (20% of the OMM proteome), whereas the rest of the 103 proteins (80% of the OMM proteome) are not GO annotated as “Mitochondria” and are therefore OMM interacting proteins that we are interested in our screening.

Column 8: Comparison of our neuronal OMM proteome with the published OMM+NES proteome and OMM exclusive proteome in HEK cells (Hung et al., 2017) reveals 69 overlapping proteins (53% of the neuronal OMM proteome).

Column 9: Actin (Actb) interactors found in the OMM proteome reveals 21 proteins.

Column 10: Tubulin (Tuba1a) interactors found in the OMM proteome reveals 6 proteins.

**Table S2. Related to Figure 2. Actin interactors found in the OMM proteome comprising 21 proteins**

Column 1: UniProt ID of the actin (Actb) interactors in the OMM proteome

Column 2: Gene names of the actin interactors in the OMM proteome

Column 3: Protein names of the actin interactors in the OMM proteome

Columns 4, 5, 6: Presence or absence of the actin interactors in the OMM proteome in replicates 1, 2, and 3, respectively.

Column 7: GO annotation of the actin interactors in the OMM proteome with the term "Mitochondria" reveals 5 proteins, whereas the rest of the 16 proteins are not GO annotated as "Mitochondria".

Column 8: 21 Actin interactors found in the OMM.

Column 9: 5 Tubulin (Tuba1a) interactors that are also OMM-actin interactors.

Column 10: Protein function as in UniProt website.

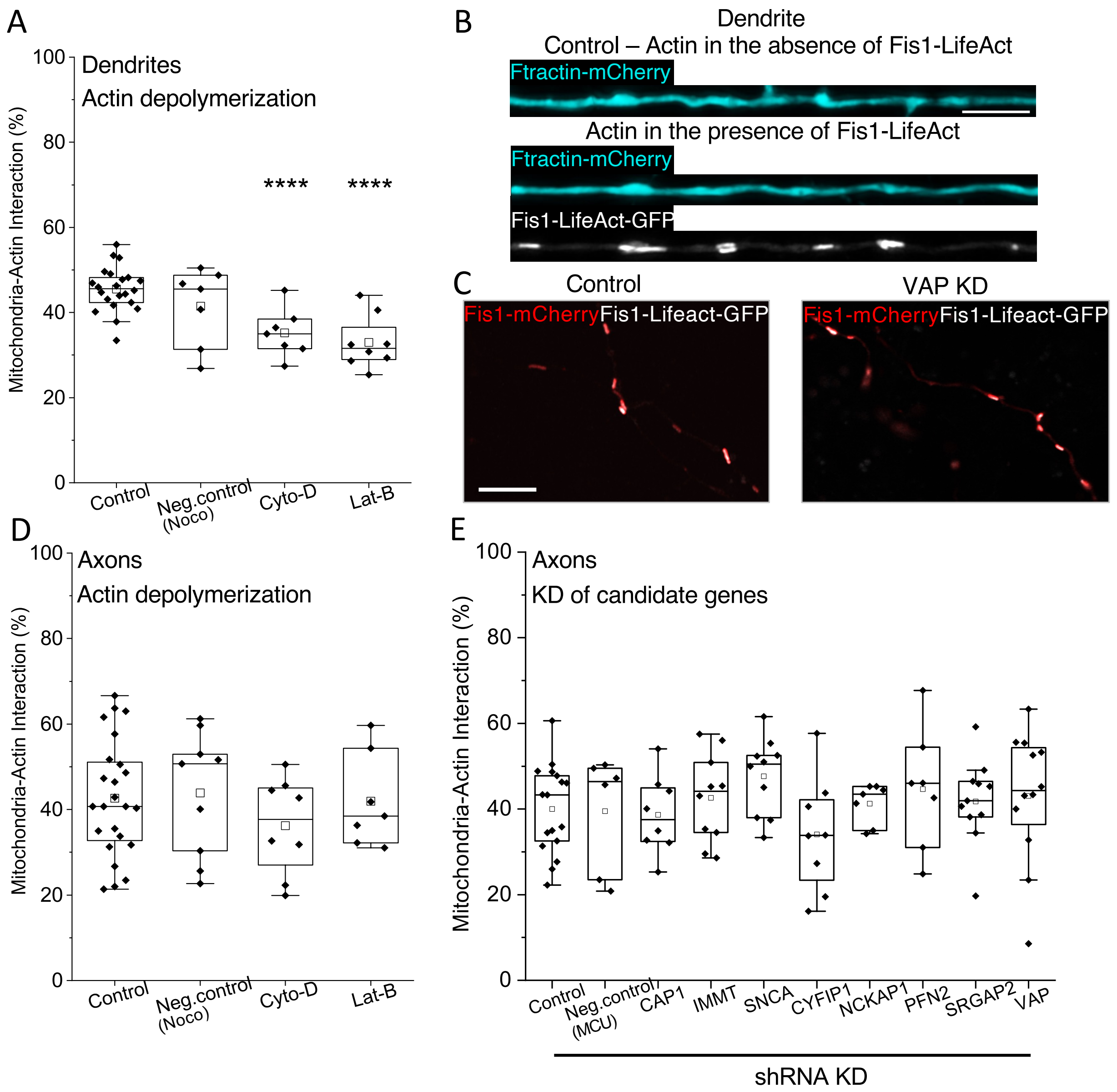


**Figure S3. Related to Figure 3. Identified candidate proteins are not required for mitochondrial-actin interaction in axons**

(A) Significant reduction in the average mitochondria-actin interaction percentage on actin depolymerization using Cytochalasin-D (Cyto-D) and Latrunculin-B (Lat-B), and not on microtubule depolymerization using Nocodazole as negative control (Neg. control, Noco), compared to Control (absence of any treatment). n $\geq$ 7 neurons from $\geq$ 2 animals. One-way ANOVA, Tukey test, ****p $\leq$ 0.00001. (B) Representative image of a neuronal dendrite transfected with Ftractin-mCherry (cyan), as an actin visualization marker (Control, Top), showed no obvious change or abnormal aggregation of actin when co-transfected with Fis1-Lifeact-GFP (Bottom, white). Scale bar, 5 μm. (C) Representative images showing a negligible difference in mitochondrial (red) regions interacting with actin (white) between control shRNA-expressing axons (left) and VAP KD axons (right). Scale bar, 5 μm. (D) The average mitochondria-actin interaction percentage measured in axons showed no significant difference on actin depolymerization using Cyto-D, Lat-B, and microtubule depolymerization using Noco, compared to Control. $n \geq 7$ neurons from $\geq 2$ animals. One-way ANOVA, Tukey test, $p = 0.6$. (E) The average mitochondria-actin interaction percentage measured in axons showed no significant difference on knocking down the 8 protein candidates of interest. Control (control shRNA), Neg. control (MCU shRNA as negative control), CAP1, IMMT, SNCA, CYFIP1, NCKAP1, PFN2, SRGAP2, VAP. $n \geq 6$ neurons from $\geq 2$ animals. One-way ANOVA, Tukey test, $p = 0.5$.

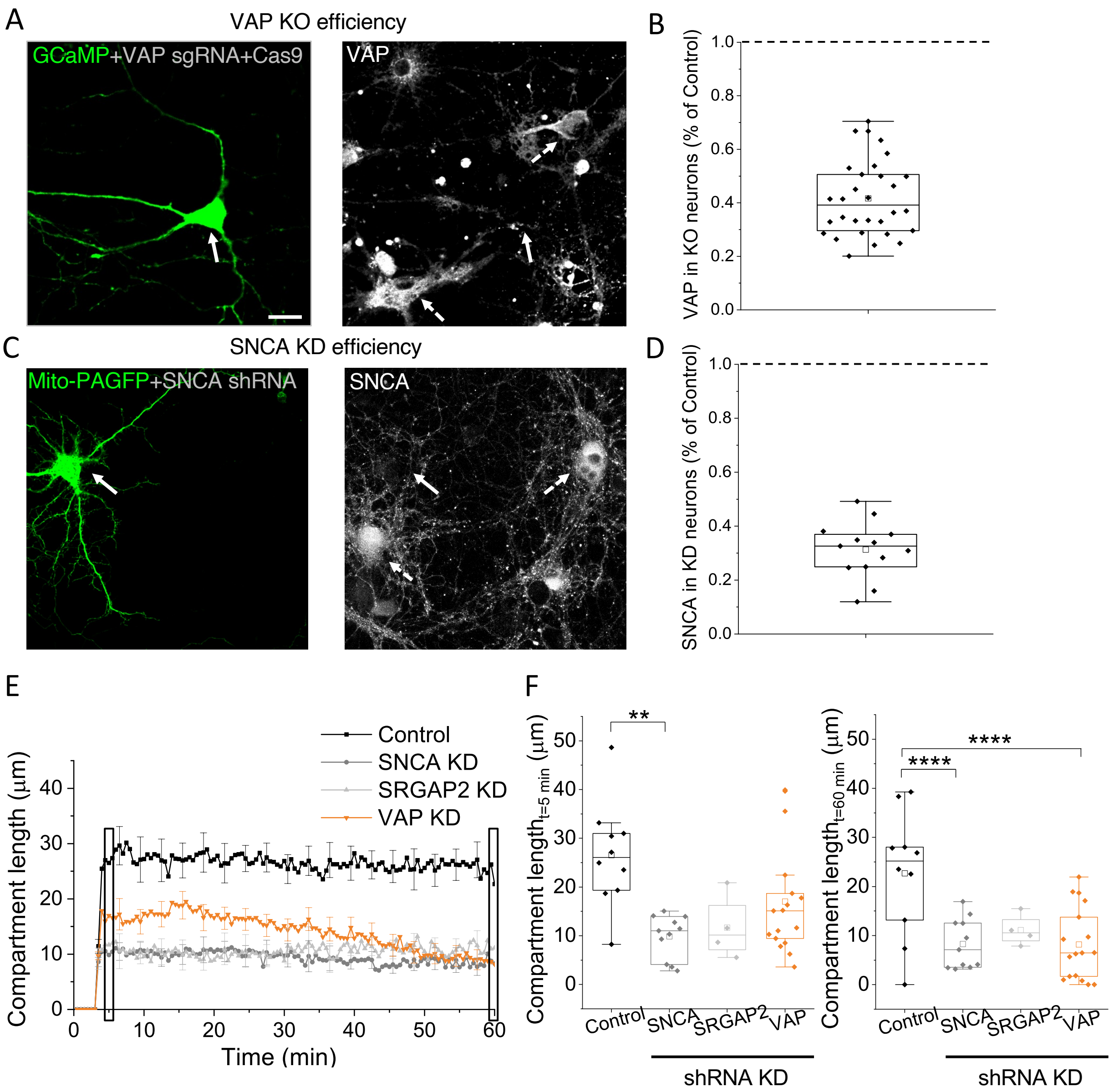


**Figure S4. Related to Figure 4. Knockdown efficiency and mitochondrial compartment length determination**

(A) Representative image of a transfected neuron (white arrow) expressing GCaMP (left, green) and VAP(A/B) CRISPR-Cas9 sgRNAs, and immunostained for VAP(A/B) (right, gray) shows reduced VAP immunofluorescence compared to adjacent, untransfected, control neurons (dashed white arrows), identified using MAP2 as a neuronal marker (image not shown). Similar results were obtained with VAP shRNA KD (data not shown). Scale bar, 20 µm. (B) Reduced average VAP immunofluorescence measured in VAP(A/B) CRISPR-Cas9 sgRNA transfected neurons compared to adjacent, untransfected control neurons. n = 30 neurons from 10 animals.

(C) Representative image of a transfected neuron (white arrow) expressing Mito-PAGFP (left, green) and SNCA shRNA, and immunostained for SNCA (right, gray) shows reduced SNCA immunofluorescence compared to adjacent, untransfected, control neurons (dashed white arrows), identified using MAP2 as a neuronal marker (image not shown). Scale bar, 20 μm. (D) Reduced average SNCA immunofluorescence measured in SNCA shRNA transfected neurons compared to adjacent, untransfected control neurons. n = 13 neurons from 2 animals. SRGAP2 knockdown efficiency could not be confirmed due to a lack of reliable antibodies. However, the successful expression of SRGAP2 shRNA in neurons was confirmed with a Td-Tomato fluorescent tag. (E) The average time course of the photoactivated mitochondrial compartment lengths show stable mitochondrial compartments in control shRNA expressing dendrites (gray) but shortened compartments in SNCA (gray), SRGAP2 KD (light gray), and VAP KD (orange) dendrites. $n \geq 4$ neurons from $\geq 2$ animals. (F) The corresponding compartment lengths from Fig. S4E (black rectangles) measured at 5 min (left) and 60 min (right) post-photoactivation show a significant reduction in mitochondrial compartment length in SNCA KD dendrites at 5 and 60 minutes post-photoactivation, and in VAP KD dendrites at 60 minutes post-photoactivation compared to those expressing control shRNA, while SRGAP2 KD dendrites did not show statistical significance. One-way ANOVA, Tukey test, $^{**}p \leq 0.01$, $^{****}p \leq 0.0001$.

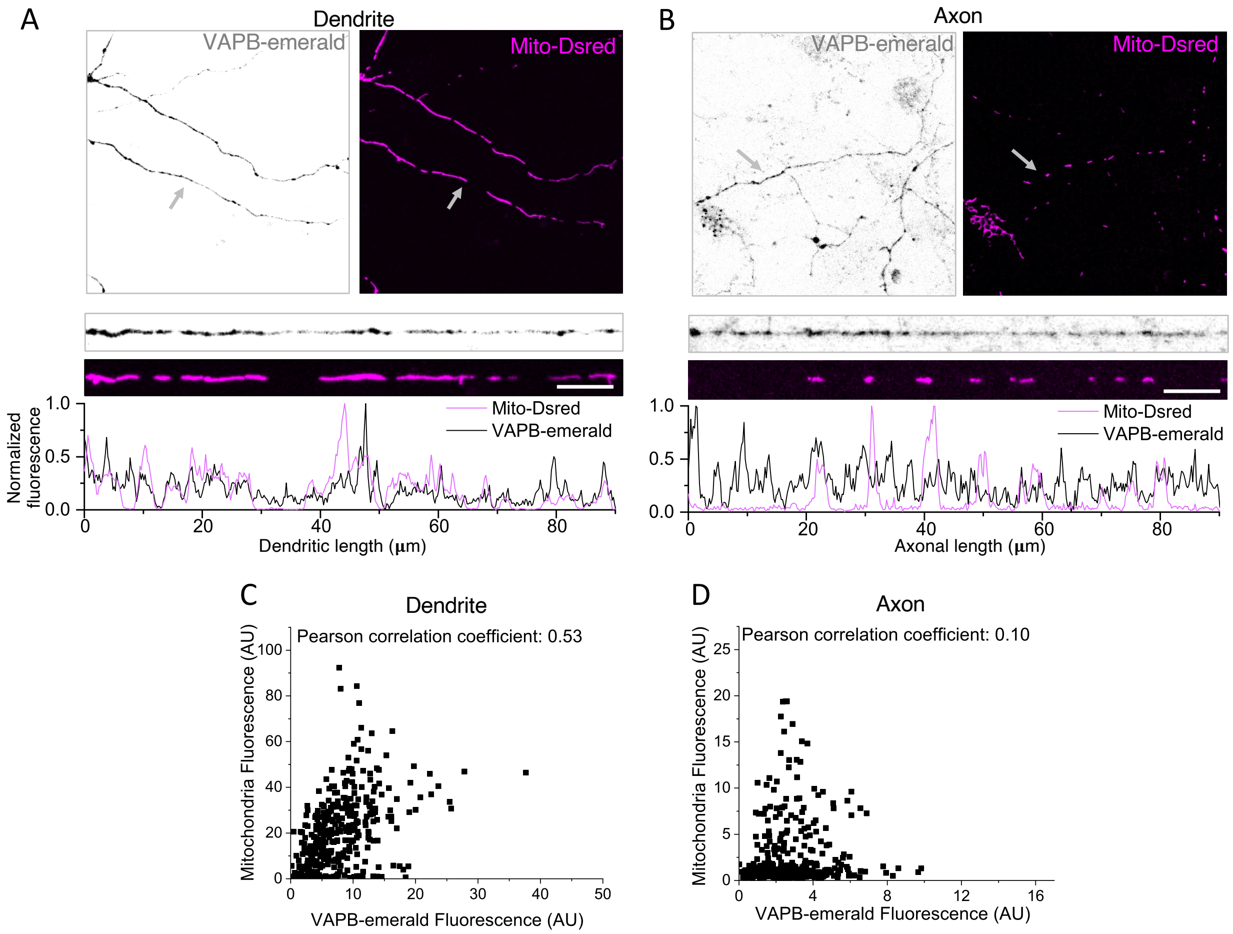


**Figure S5. Related to Figure 5. VAPB is enriched near dendritic mitochondria but not near axonal mitochondria in live neurons**

(A) Representative images of dendritic compartments from live neurons expressing VAPB (left, VAPB-emerald, gray) and mitochondria (right, Mito-DsRed, magenta). Gray arrows depict dendrites magnified for better visualization (bottom insets), showing local enrichment of VAPB near dendritic mitochondria in live neurons. Scale bar, 5 μm. (B) Representative images of axonal compartments from live neurons expressing VAPB (left, VAPB-emerald, gray) and mitochondria (right, Mito-DsRed, magenta). Gray arrows depict axons magnified for better visualization (bottom insets), showing negligible enrichment of VAPB near axonal mitochondria in live neurons. Scale bar, 5 μm. Representative correlation between individual fluorescent pixel intensities of mitochondria (Mito-DsRed) and VAPB (VAPB-emerald) show a strong correlation in the dendrite (C) but not in the axon (D). Summarized data is available in Figure 5D.

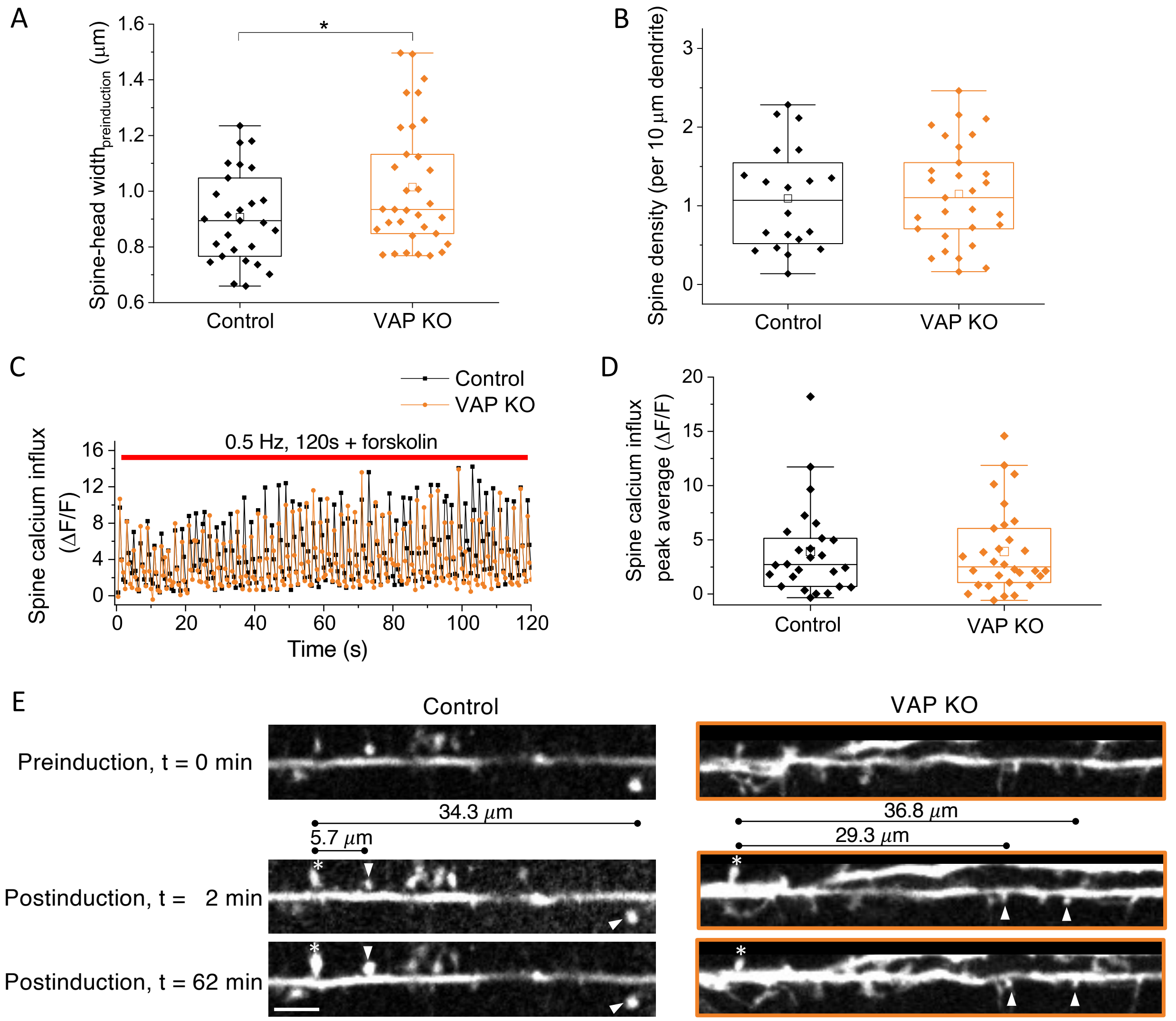


**Figure S6. Related to Figure 6. VAP depletion does not affect spine size, density, and calcium influx**

(A) Spine-head width showed a modest and negligible increase in VAP KO neurons compared to control sgRNA-expressing neurons at baseline. $n \geq 27$ spines from $\geq 7$ animals. Two sample t-Test, $*p \leq 0.05$. (B) Spine density was not affected in VAP KO neurons compared with control sgRNA-expressing neurons at baseline. $n \geq 20$ dendrites from $\geq 5$ animals. Two sample t-Test, $p = 0.8$. Representative trace (C) and peak average (D) of the spine calcium influx was not affected in VAP KO neurons, compared with control sgRNA-expressing neurons, during synaptic plasticity induction. $n \geq 26$ spines from $\geq 7$ animals. Two sample t-Test, $p = 1.0$. (E) Representative plasticity-induced spine (white asterisk) measured at 0 min (preinduction), 2 min (postinduction), and 62 min (postinduction) showed an increase in spine-head width in plasticity-induced spines and adjacent, PSD95 or Homer2 positive, unstimulated spines (white arrowhead) at 5.7 μm distance (less than 30 μm) but not in spines at 34.3 μm distance (beyond 30 μm), in neurons expressing control sgRNA (Left, Control). VAP KO neurons exhibited structural plasticity in the plasticity-induced spine 2 min postinduction but not 62 min postinduction (white asterisk) (Right, VAP KO, orange box). VAP KO neurons also did not exhibit structural plasticity in adjacent, PSD95 or Homer2 positive, unstimulated spines (white arrowhead) at 29.3 μm distance (less than 30 μm) and in spines at 36.8 μm distance (beyond 30 μm) (Right, VAP KO, orange box). Scale bar, 5 μm.
